# Global Environmental Genomics Reveals Vast Uncultivated Protist Diversity and Taxonomic Blind Spots

**DOI:** 10.1101/2025.03.26.645542

**Authors:** Miguel F. Romero-Gutiérrez, Arianna I. Krinos, Xyrus Maurer-Alcala, John A. Burns, Ramunas Stepanauskas, Tanja Woyke, Frederik Schulz

## Abstract

Protists, defined as eukaryotes distinct from animals, plants, and fungi, are a polyphyletic group that dominates the eukaryotic tree of life, exhibiting significant phylogenetic diversity and fulfilling critical ecological roles. Historically, research has prioritized protists associated with animals and plants, particularly those of medical significance, thereby overlooking the majority of protist diversity. Conventional molecular methods, such as 18S rRNA gene amplicon sequencing, frequently encounter limitations, including primer binding bias and PCR bias caused by gene length variations, resulting in a biased representation of protistan diversity. Further, most protist lineages are notoriously difficult to cultivate. Here, we apply a cultivation-independent approach in which we analyzed over 30,000 assembled metagenomes and protist single cell genomes and 21 long 18S rRNA gene amplicon data sets from various global ecosystems, including marine, freshwater, and soil environments. We recovered 157,956 18S rRNA gene sequences (≥800 bp), which clustered into 103,338 operational taxonomic units (OTUs) at 97% sequence identity and 24,438 OTUs at 85% identity. Notably, 81% of 9,543 non-singleton clusters at 85% identity classified as protists consisted exclusively of environmental sequences, uncovering a wealth of novel, uncultivated, and unclassified protist diversity. A comprehensive taxonomic framework of eukaryotes based on concatenated 18S and 28S rRNA genes that incorporated most novel lineages revealed substantial underrepresentation of Amoebozoa, Discoba, and Rhizaria in reference databases, with many lacking isolate or genome sequence representation. Further, we identified 13 eukaryotic lineages with novelty on higher taxonomic ranks, such as class and phylum-level, that lack representation in public databases. The corresponding 85% OTUs were primarily affiliated with Excavata, with some branching deeply in the eukaryotic tree. Comprehensive analysis of the global distribution of protists revealed uneven microbial diversity across supergroups and ecosystems, with notable novelty particularly in soil and marine environments. We then examined co-occurrence between protists and prokaryotes, predicting putative symbiotic or predator-prey relationships, particularly among understudied protist groups with bacteria such as Verrucomicrobia and Rickettsiales. Our results substantially enhance the understanding of protistan diversity and distribution, revealing taxonomic blind spots and laying groundwork for future studies of these organisms’ ecological roles.

## Introduction

The vast majority of eukaryotic diversity lies outside the well-established kingdoms of Animalia, Fungi, and Plantae. These organisms have traditionally been referred to collectively as ‘protists’, a term that encompasses a highly diverse and non-monophyletic assemblage of lineages. In addition to their remarkable diversity, protists exhibit a wide array of cellular structures and lifestyles (Foissner 2006; Adl et al. 2007). Previous research has largely focused on protists associated with animals and plants, particularly those of medical relevance (Collins and Jeffery 2007; Horn 2022; Einarsson et al. 2016). Well-characterized lineages include the TSAR group, which comprises Stramenopiles (e.g., diatoms and oomycetes), Alveolata (e.g., ciliates, dinoflagellates, and apicomplexan parasites), Rhizaria (e.g., foraminifera and radiolarians), as well as the Amoebozoa supergroup (sensu stricto amoebae) and the Discoba division (which includes heteroloboseans and euglenoids, including kinetoplastids). Even less well studied eukaryotic lineages include Metamonada, which encompass instances of symbiotic or parasitic groups like diplomonads and parabasalids (Céza et al. 2022). Recently, advances in molecular methods, such as cultivation-independent single-cell sequencing (Woyke et al. 2017), have facilitated the exploration of phylogenetic associations among selected uncultivated groups, resulting in the identification of new eukaryotic supergroups, such as Hemimastigophora (Foissner et al. 1988; Lax et al. 2018) and Provora (Tikhonenkov et al. 2022), with additional members continually being added as interest in these novel lineages expands (Eglit et al. 2024).

Molecular methods such as 18S rRNA gene amplicon sequencing have proven invaluable for assessing eukaryotic diversity in various environmental contexts. In oceanic plankton communities, studies revealed that heterotrophic protists represent the most diverse group, despite metazoans dominating in terms of abundance (de Vargas et al. 2015). A protist-centric survey in select soil environments indicated that Cercozoa are the most prevalent group, revealing correlations between protistan and bacterial taxa, including evidence of potential predator-prey interactions (Oliverio et al. 2020). At a global level, soil eukaryotic diversity has primarily been evaluated with respect to specific groups such as Fungi and testate amoebae (Fernández et al. 2016; Bahram et al. 2018). Diversity of eukaryotes in soil typically exhibits higher values in the Simpson diversity index in equatorial regions and intermediate latitudes, indicating that diversity peaks in temperate climates, likely linked to water-energy balance and present similar patterns such as plants and animals (Fernández et al. 2016; Bahram et al. 2018). A comprehensive meta-analysis that incorporated data from marine, freshwater, and topsoil ecosystems demonstrated that topsoil harbors greater protistan diversity compared to aquatic environments (Singer et al. 2021). Despite these extensive studies on eukaryotic diversity and community structure, recent findings indicate that an overwhelming majority of protistan taxa remain undocumented. For example, an analysis suggested that one-third of operational taxonomic groups (OTUs) could not be assigned to known eukaryotic groups (de Vargas et al. 2015). Similarly, new lineages of protists, at both the phylum and supergroup levels, including Hemimastigophora and Provora, have only recently been described (Foissner et al. 1988; Lax et al. 2018; Tikhonenkov et al. 2022).

The 18S rRNA gene (also referred to as 18S rDNA) has emerged as the most frequently used marker gene for identifying eukaryotic organisms in environmental studies. This gene encodes the small subunit of the ribosomal RNA and is universally present in all nuclear genomes, comprising regions of both high conservation and high variability (Wuyts et al. 2001). Public databases, such as Silva (Quast et al. 2013) and the Protist Ribosomal Reference (PR2; Guillou et al. 2013), have served as repositories for these sequences for decades. Polymerase chain reaction (PCR)-based approaches (metabarcoding) are typically used to recover this marker gene with high sensitivity in environmental samples. However, due to the constraints of short-read DNA sequencing technologies, only a limited portion of the 18S rDNA is typically screened, predominantly the V4 to V6 and V9 regions (Hugerth et al. 2014; de Vargas et al. 2015; Oliverio et al. 2020). Unlike its prokaryotic counterpart (the 16S rRNA gene), the 18S rRNA gene exhibits pronounced length variation, ranging from a minimum of 1.5 kb (in Diplomonadida) to a maximum of 6.4 kb (in Amoebozoa), with the mature 18S rRNA molecule potentially reaching lengths of up to 2.3 kb (Xie et al. 2011). This variability can be attributed to length-variable regions that frequently contain introns, some exceeding 1 kb, especially in regions commonly amplified (Xie et al. 2011). Such disparities in length, coupled with the rapid evolution of the 18S rRNA gene in certain lineages (Meyer et al. 2010), may bias PCR-based metabarcoding studies, as only a fraction of the eukaryotic diversity in the environment can be captured. Recently, long amplicon sequencing has been deployed to recover multiple markers from the rRNA gene locus within eukaryotic genomes (Jamy et al. 2022). Although this technique solves some of the challenges associated with conventional methods (Overgaard et al. 2024), primer binding bias remains a concern due to sequence divergence.

An alternative approach that is able to further mitigate limitations of amplicon-based methods is shotgun metagenomics as it does not employ specific primers for amplifying target sequences. Shotgun metagenomics-based screening involves analyzing assembled sequences from the genomes of all taxa present in the sample for rRNA genes. Depending on sequencing depth, genomic information from rarer taxa can also be retrieved. Targeted marker gene recovery from metagenomic assemblies is more sensitive than genome-resolved metagenomics, which relies on successful binning. This gene-centric approach provides higher sensitivity by enabling the analysis of contigs from low-abundance organisms or fragmented genomes that fall below the thresholds required for accurate binning (Nayfach et al. 2021). However, this method remains susceptible to artifacts inherent to the assembly process itself, including sequence fragmentation, chimerism, and the collapse of strain-level diversity into consensus sequences (Sczyrba et al. 2017). In our study, we employed a shotgun metagenomics approach combined with marker gene extraction from assembled contigs. We leverage over 30,000 assembled metagenomes to conduct a global census of eukaryotic diversity. Given the potential confounders introduced by metagenomic sequence assembly, which can obscure fine-grained phylogenetic signals, we focused our analysis primarily on higher taxonomic ranks. Our primary objective was to elucidate the phylogenetic and ecosystem diversity of microbial eukaryotes on a global scale, thereby advancing the understanding of protistan diversity and distribution, including the characterization of previously unidentified lineages.

## Results and discussion

### Expansion of Known Protist Diversity

The screening of 32,520 assembled metagenomes yielded 157,956 18S rRNA genes from 18,445 sequencing datasets, all with a minimum length of 800 bp (a summary of the methods can be found in Supplementary Figure 1). Subsequently, we performed two-step clustering of these metagenome-derived sequences in conjunction with over 250,000 sequences from the curated rRNA gene databases Silva and PR2 (Quast et al. 2013; Guillou et al. 2013) and additional recent studies (Jamy et al. 2022; Tikhonenkov et al. 2022), yielding a total of 103,338 sequence clusters (OTUs) at 97% sequence identity and an additional 24,438 OTUs at 85% identity to account for the high variability of correspondence between sequence identity of the 18S rRNA gene and taxonomic ranks. We discarded all 11,200 85% identity clusters containing only a single sequence to remove potential chimeric sequences, retaining 13,238 clusters with two or more sequences for further analysis. Further, we excluded from our dataset sequences classified within the Streptophyta_X, Fungi, and Metazoa subdivisions and chimeric representative 18S gene sequences to focus our analyses on protistan diversity. The filtered dataset presented 112,768 metagenome-derived 18S sequences clustered within 9,543 clusters at 85% sequence identity. As reference databases such as Silva and PR2 include sequences originating from both cultured isolates as well as environmental sources, we were able to estimate the proportion of sequences corresponding to uncultivated microbes within our dataset.

Our analysis revealed that 81% of the 85% identity clusters (n=7,716) were exclusively comprised of environmental sequences, while 19% (n=1,780) included sequences from isolates and less than 1% (n=47) contained sequences derived from genomes. Clusters that contained sequences from isolates (18S rRNA genes) were generally larger, whereas the metagenomic-only clusters tended to be smaller (Figure 1A). Clusters containing genome sequences had an average of 1,031.64 members, greatly exceeding the average of 54.52 members for clusters containing sequences from isolates. Notably, the mean size of clusters containing solely environmental sequences was calculated at 10.39 (SD=24.04). Among the largest clusters containing only sequences from reference databases were sequences from Piroplasmorida (Apicomplexa, size=1,220) and Cryptosporida (Apicomplexa, size=910). Conversely, the largest clusters including metagenomic sequences comprised sequences from Syndiniales (Dinoflagellata, size=1,028) and Fornicata_X (Fornicata, size=487). The largest cluster incorporating genome sequences was affiliated with Colpodellida (Protalveolata, size=10,291), while the largest cluster containing isolate-derived sequences without genome information was ascribed to Syndiniales Group II (Syndiniales, size=4,275). Notably, the largest cluster exclusively composed of environmental sequences was also designated to Syndiniales Group II (Syndiniales, size=1,028). To further investigate the clusters with only reference sequences, we estimated the number of distinct taxa found per cluster.

**Figure 1.**
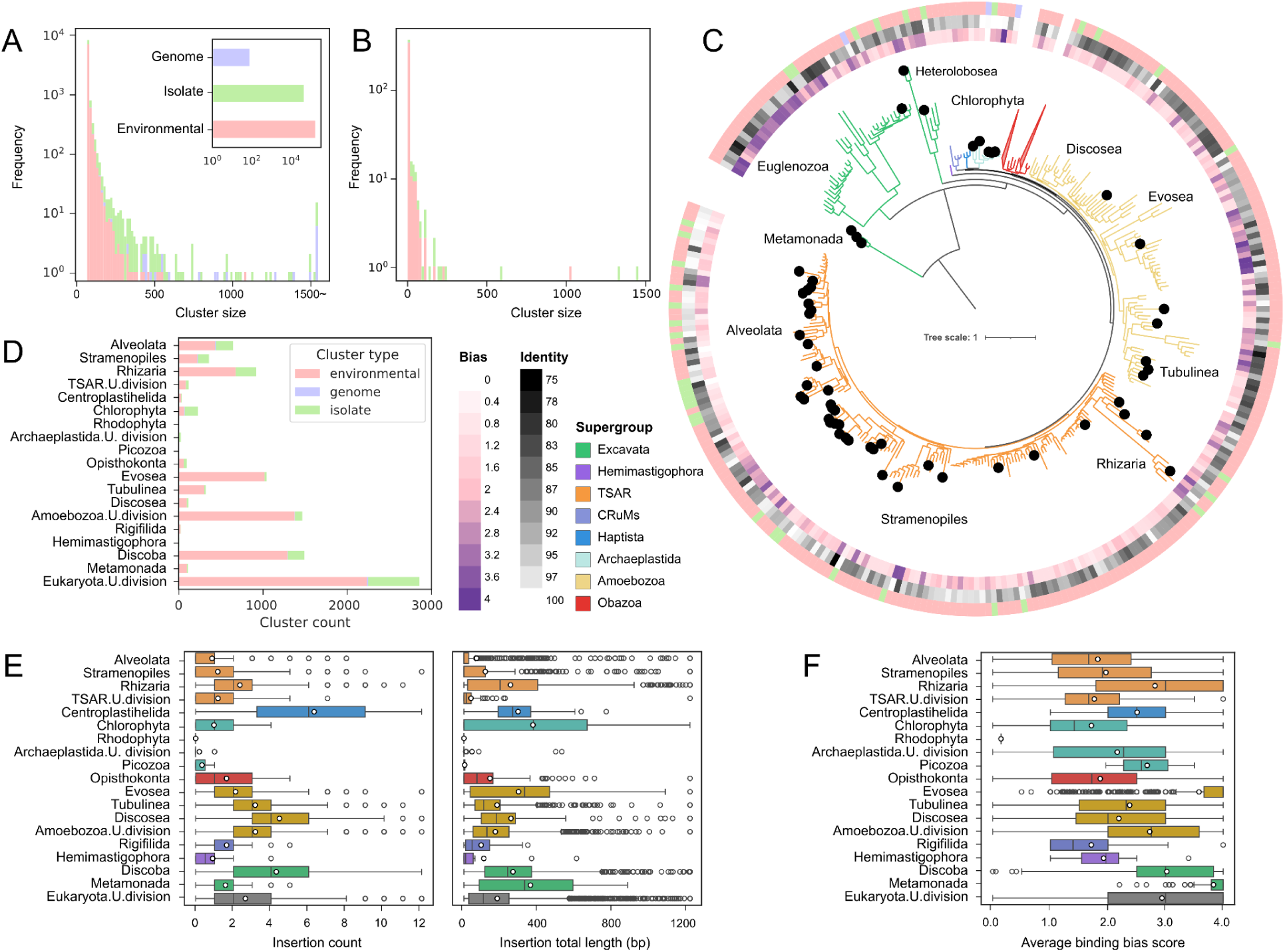
Phylogenetic Diversity of Eukaryotes. (A) Histogram illustrating the distribution of cluster sizes, color-coded based on the origin of the included sequences. The inset represents the number of clustered sequences from each source. Data corresponds to 85% identity clusters after discarding singletons, chimeras, and non-protists. (B) Cluster size distribution of the OTUs that contained both 18S and 28S rRNA genes, which were employed to construct the eukaryotic reference tree. (C) Reference tree inferred from a concatenated alignment of the 18S and 28S rRNA genes derived from OTUs that had representative members with both genes present. The heat map on the innermost track indicates the cluster-wide average of the accumulated primer binding bias score of the corresponding 18S rRNA gene sequences utilizing a set of four commonly used general eukaryotic primer pairs. A score of 4 (dark purple) suggests that the sequence would not bind to any of the primer pairs, while a score of 0 (white) implies it would bind to all of them. The adjacent heatmap delineates sequence identity compared to the best hit within our combined 18S rRNA gene sequence database. The outermost track denotes the cluster composition corresponding to the legend illustrated in panel A. Black circles at the tip of the leaves indicate that the respective clusters are composed exclusively by environmental sequences extracted in this study. Clades belonging to Fungi, Metazoa, and Streptophyta are collapsed. (D) Relative abundance of clusters for each protist division within the analyzed environmental samples. (E) Distribution of insertion count and cumulative insertion length among protist divisions. Each point shows the corresponding data for each OTU representative sequence. Insertions are defined as sequences present in each 18S rRNA gene sequence not represented in the reference covariance model. Data of insertions with length >= 10 bp is shown. Box plots depict the interquartile range (IQR, 25th to 75th percentiles), with horizontal lines representing median values. Whiskers extend to the most extreme data points within 1.5 × IQR from the box boundaries, and data points beyond this range are displayed as individual circles, representing statistical outliers. White circles indicate mean values. (F) Distribution of predicted primer binding bias score for each OTU representative sequence ordered by its taxonomic classification at the division level. Box plots show the interquartile range with median values (horizontal lines). Whiskers extend to data points within 1.5 × IQR, with outliers represented as individual circles. Mean values are shown as white circles.

Our data-driven approach generated clusters containing, on average, 1.5 genera (SD=1.61), 1.29 families (SD=0.92), and 1.13 orders (SD=0.51) (Supplementary Figure 2). Similarly, when considering only well-resolved orders (excluding unclassified .U.order and unresolved _X lineages), each order encompassed an average of 1.29 families (SD=0.93) and 1.50 genera (SD=1.63). These data suggest that the 85% sequence identity threshold used for clustering corresponds approximately to the taxonomic level of order or family. Although metagenomic assembly is susceptible to inherent artifacts including sequence fragmentation, chimerism, the collapse of strain-level diversity into consensus sequences (Sczyrba et al. 2017), and exhibits low sensitivity for low-abundance members, we mitigated these issues by focusing our analyses on higher taxonomic ranks.

Our results indicate that protists recognized as parasites of animals and humans, such as Piroplasmorida (Apicomplexa, 1,220 sequences) and Cryptosporida (Apicomplexa, 910 sequences), are the predominant representatives within reference databases, revealing a pronounced sampling bias in the representation of host-associated microbial eukaryotes (del Campo et al. 2014). Some parasitic protists, such as apicomplexans, were also found to be dominant in soil microbiomes (Mahé et al. 2017). Consequently, it is unsurprising to encounter sizable clusters of sequences derived from parasitic protists, whether from isolates or environmental samples, with the latter exhibiting a more balanced representation of the diversity of these organisms.

To enhance the taxonomic annotation of sequences and to better reflect the phylogenetic relationships among eukaryotes, we constructed a eukaryotic reference tree from 85% OTU cluster representatives that also carried a 28S rRNA gene on the same assembled contig (1,438 candidate OTUs; 577 retained in the final tree after length filtering and rogue-taxon removal, see Methods). This subset of our dataset excluded most of the highly populated clusters (Figure 1B). The resulting tree broadly aligned with the established phylogeny of eukaryotes, as it effectively grouped the Amorphea together, distinctly separate from the Diaphoretickes, in agreement with observations from both rRNA and protein-based alignments (Jamy et al. 2022; Torruella et al. 2024). Further, Metamonads were positioned near the root, consistent with phylogenomic analysis of the root of the eukaryotic tree (Al Jewari and Baldauf 2023; Williamson et al. 2025) . Due to high mutation rates and pronounced length variability, certain groups such as Microsporidia (Vossbrinck et al. 1987) were excluded from the reference tree, leaving their representatives unclassified within our phylogenetic taxonomic annotation. Numerous clades grouped within Metamonada, Stramenopiles, Alveolata, Amoebozoa, Metazoa, and Fungi divisions entirely lacked sequences derived from isolates or genomic sources (Figure 1C). Clusters associated with these clades also tended to exhibit representative sequences with low blast alignment identity to reference sequences and increased average predicted primer binding bias (Figure 1C) compared to sequences from other clusters, suggesting these represent taxonomic blindspots overlooked in prior PCR amplification-based studies (Morey et al. 2024).

Our census of protistan diversity revealed an uneven taxon richness across various supergroups, with Amoebozoa (n=2,874), and TSAR (n=1,887) exhibiting the highest number of OTUs. At the division level, Discoba harbored the largest diversity, predominantly characterized by sequences derived from the environment. The divisions classified under Amoebozoa exhibited a high proportion of clusters exclusive to environmental samples (87%), most of which remained unclassified at the division level (n=1,423) (Figure 1D). Similarly, Discoba, the second most diverse division, demonstrated novelty in sequences presented as a substantially larger proportion of clusters derived exclusively from metagenomic sequences (n=1,264), with only 200 clusters containing sequences from isolates and a mere 6 clusters encompassing genome-derived sequences. In the context of the TSAR supergroup, Rhizaria emerged as the most diverse division, with 72% of the 877 clusters comprising entirely environmental sequences. Conversely, Amoebozoa’s high diversity demonstrates its adaptive radiation across soil and aquatic habitats through varied locomotion mechanisms and predatory strategies (Tekle et al. 2022), which could be true also for some groups of discobans. Within the TSAR supergroup, the dominance of Rhizaria can be seen as a product of their rapid evolutionary rates (Burki and Pawlowski 2006). The resulting diversification, which is particularly well-documented in marine systems, has resulted in highly divergent SSU rRNA gene sequences even among closely related taxa (Schuler et al. 2018). The 85% sequence identity threshold employed for our clustering was specifically chosen to accommodate this substantial sequence divergence. This approach successfully resolved numerous diverse rhizarian lineages, underscoring the degree to which current cultivation-based methods fail to capture the true phylogenetic breadth of these major eukaryotic groups.

Representative sequences that could not be assigned to established eukaryotic supergroups were designated ‘Eukaryota.U’. Consequently, ‘Eukaryota.U’ does not represent a single monophyletic group; rather, it is a polyphyletic assemblage of multiple, taxonomically distinct, and unclassified eukaryotic lineages. Among these, 2,198 (78%) were exclusively derived from environmental sequences, while 612 clusters contained sequences from isolates and 13 clusters harbored sequences from genomes. Among the reference-only clusters that were unassigned to any supergroup using the tree-placement method, most were represented in the reference tree according to sequence annotation, such as Cnidaria (129 clusters), or had a sister group present within the same supergroup or division, exemplified by Microsporidia (36 clusters). This finding is somewhat expected for lineages such as Microsporidia, which are characterized by short yet highly divergent 18S rRNA gene sequences (Vossbrinck et al. 1987). Challenges in supergroup assignment using tree-placement methods often stem from incomplete reference databases or rapid evolutionary rates. Lineages with rapid sequence evolution or sparse reference coverage often resolve at lower ranks but not at the supergroup level, as previously reported (Berger et al. 2011).

We found differences in insertion number and length among eukaryotic supergroups, with Centroplasthelida with the highest mean number of insertions (6.71, SD=3.51), and Chlorophyta with the longest mean insertion size (688.70, SD=393.80; Figure 1E). Even with variation within lineages of eukaryotic supergroups (Xie et al. 2011), we found differences between groups. Predicted binding bias scores also varied among groups, with those with the highest proportion of phylogenetic novelty (Figure 1D) showing also a high mean binding bias score, such as Metamonada (mean score=3.83, SD=0.33), Evosea (3.59, SD=0.81), Discoba (3.03), and Eukaryota.U.division (unclassified at the supergroup level, mean score=2.94, SD=1.04). This observation could explain why some lineages within these groups might have remained as taxonomic blindspots in metabarcoding studies.

On average, each of the 12,398 metagenomic sequencing datasets provided 9.10 (SD=24) 18S sequences to our analysis. This low per-sample yield likely reflects a combination of sampling and library preparation commonly optimized for prokaryotic biomass, low relative abundance and patchy distribution of microbial eukaryotes in many metagenomes, and assembly-related challenges for rDNA loci (e.g., introns, repeats, strain heterogeneity, and fragmentation), compounded by our conservative length threshold chosen to support robust phylogenetic placement. This is a common limitation of whole metagenome shotgun sequencing analyses that aim to recover eukaryotic sequences. For instance, bacterial MAGs are more frequently recovered than eukaryotic MAGs (Bowers et al. 2025; Duncan et al. 2022). We therefore focus on higher-rank patterns as candidates for targeted validation.

### The Global Distribution of Protists

The majority of 18S rRNA gene sequences recovered (present in non-singleton clusters) were from the Integrated Microbial Genome/Metagenome (IMG/M) database (Chen et al. 2023), representing samples from all continents, encompassing longitudes and latitudes from pole to pole (Figure 2A). The main ecosystem type sampled presenting 18S genes was marine (n=3,866), followed by freshwater (n=2,888) and soil (n=2,530). Here, “soil” refers to the GOLD ecosystem type annotation in IMG/M and encompasses a broad set of soil-associated terrestrial subtypes (e.g., peatlands/wetlands, permafrost/glacial soils, organic horizons and bulk soils); therefore, “soil” patterns should not be interpreted as applying exclusively to classical mineral topsoils. Among the frequently observed subtypes, lake samples (n=1,469), oceanic (n=884), and peat samples (n=841) were predominant (Supplementary Table 1). In contrast, datasets without any identified 18S rRNA gene sequences were most frequently associated with subtypes such as subway (n=1,028), phylloplane/leaf (n=393), and Coalbed methane well (n=317) (Supplementary Table 2), the presence of 16S rRNA genes in the same samples indicates that these may represent a sample preparation bias or a harsher environment for growth of microbial eukaryotes. Despite the broad spectrum from which these samples were gathered, the majority originated from latitudes ranging from 60° to 30° North, particularly within North America and Central Europe (Figure 2B). Key factors influencing protist diversity include water availability, pH, and nutrient availability (Zhao et al. 2023). The captured protistan diversity in freshwater and peat bogs is in agreement with what has been shown in previous studies (Lara et al. 2011; Krinos et al. 2024; Oliverio et al. 2020). In contrast, the absence of 18S rRNA gene sequences in habitats like coalbeds and built environments like subways, that are characterized by low water availability and low nutrient concentrations, reflect challenging growth conditions for eukaryotes.

**Figure 2.**
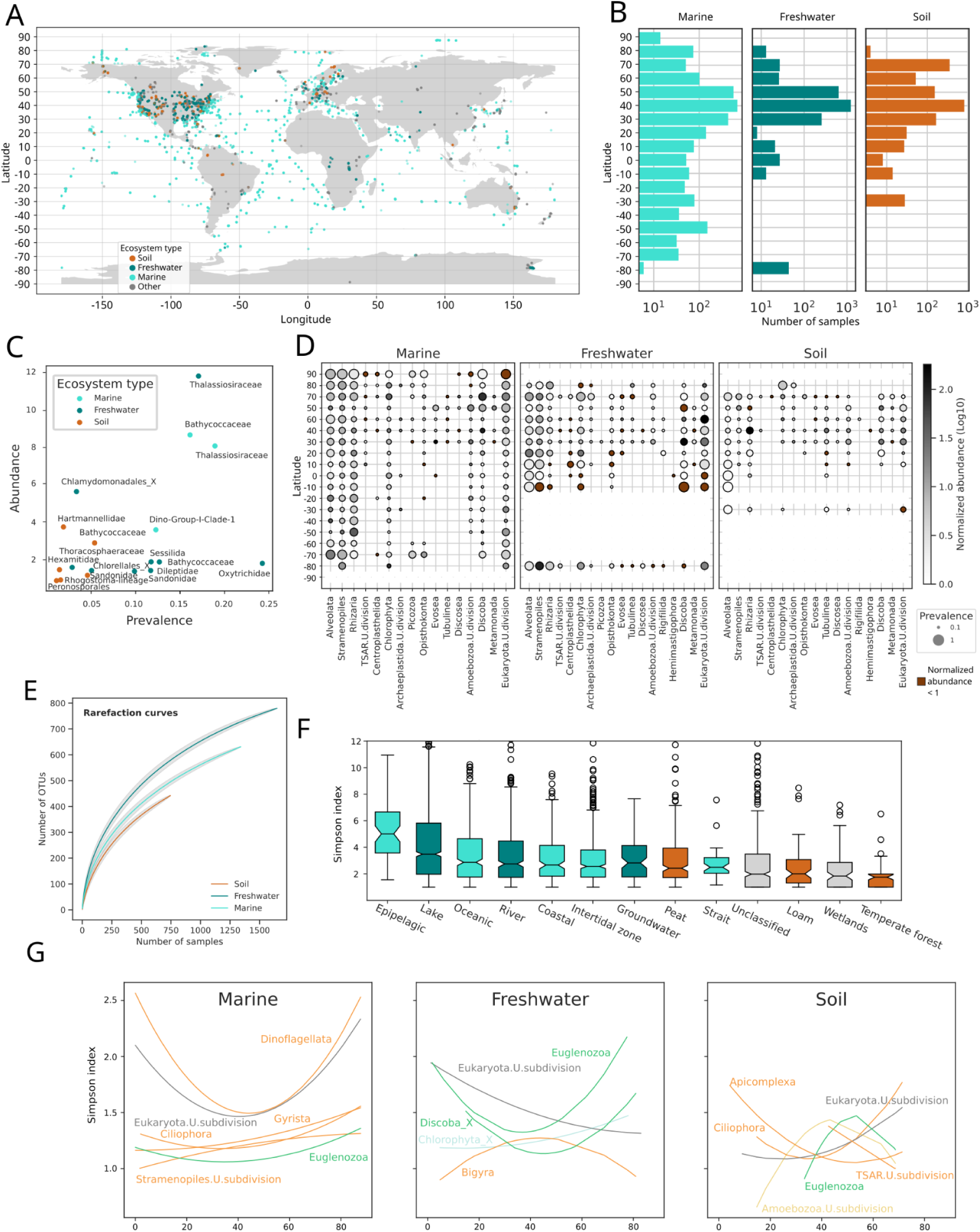
Global distribution of protists. A) World map illustrating the origin of the 11,652 samples from which 18S rRNA genes were recovered and geolocalization data is available. B) Total number of samples within each latitude range, partitioned by ecosystem type. C) Mean values of prevalence and normalized abundance for eukaryotic families that exhibited abundance and prevalence above the 80%\ quantile. Families belonging to Metazoa, Fungi, and Streptophyta have been excluded. D) Mean abundance and prevalence of each eukaryotic division across latitude ranges. Rows marked with cross marks indicate no observed samples at that latitude and ecosystem type. Brown circles indicate normalized abundance lower than 1. E) Rarefaction curves representing the mean number of observed OTUs following 10,000 randomizations. Gray shades represent standard deviation values. F) Per-sample diversity distribution for each ecosystem type with at least a 1% share of the datasets. Box plots display medians (horizontal lines), interquartile ranges (boxes), and data distribution (whiskers), with outliers shown as circles. G) Second-order polynomial fits for Simpson index values as a function of absolute latitude across different ecosystem types; only significant regressions (adjusted p-value < 0.05) are portrayed. For all panels the ‘Soil’ category follows GOLD ecosystem type annotations and includes peatland/wetland and glacial/permafrost-associated subtypes (Supplementary Table 1).

Next, we investigated the overall distribution of predominant and prevalent microbial protistan orders and families. Relative abundance was defined as the average of the normalized mean sequencing depth across all contigs pertaining to each taxonomic group, while prevalence was defined as the proportion of samples from comparable environments and latitudes in which each taxonomic group was identified. The most abundant group detected in marine ecosystems consisted of green algae from the family Bathycoccaceae (relative abundance=8.65, SD=0.78; Figure 2C), followed by diatoms (family Thalassiosiraceae; relative abundance=8.06, SD=5.92). Additionally, parasitic dinoflagellates from Ichthyodinida Clade-1 and Dino-Group-II-Clade-6 were frequently observed with relatively high abundances (relative abundance=3.64, SD=4.78 and 2.80, SD=2.30, respectively). In freshwater systems, the most abundant groups included Thalassiosiraceae (Gyrista; relative abundance=11.82, SD=37.1) and Chlamydomonadales_X (relative abundance=6.32, SD=16.72), with ciliates from the Oxytrichidae family being present across the largest number of freshwater samples (prevalence=0.24, SD=0.28). Within the GOLD ‘Soil’ category, Hartmannellidae (Tubulinea) showed the highest mean relative abundance (3.71, SD=9.37), followed by Bathycoccaceae (Chlorophyta; 2.85, SD=4.64), Hexamitidae (Fornicata; 1.41, SD=3.66), and Sandonidae (Cercozoa; 1.13, SD=1.41) (Figure 2C). Historically, diatoms and metazoans have been classified as dominant groups within marine environments, alongside chlorophytes and parasitic dinoflagellates (de Vargas et al. 2015). Free-living amoebae, such as those classified within Hartmanellidae, have been found prevalent in solid matrices, with some members found with a prevalence as high as 92% (Chauque et al. 2023). Similarly, prior studies have indicated that lineages of Cercozoa hold dominant positions within soil ecosystems, along with other groups of chlorophytes (Oliverio et al. 2020). The high Bathycoccaceae signal within the GOLD ‘Soil’ category is driven primarily by peatland/wetland and glacial/permafrost-associated subtypes (Supplementary Table 3), rather than mineral topsoils *sensu stricto*. This is consistent with water-saturated terrestrial habitats that can contain substantial algal biomass (Solari et al. 2018) and contrasts to topsoils, which contain algae from various groups, such as Chlorophyta and Bacillariophyceae (Zancan et al. 2006). This trend could also explain the abundance of the parasitic Hexamitidae that show increased prevalence with higher water availability (Wang et al. 2023). The observed differences in abundance between our findings and previous amplicon-based studies (Mahé et al. 2017; Oliverio et al. 2020) may stem from the heterogeneity of samples, enhanced gene recovery of high-abundance eukaryotes in shotgun metagenomes, and the absence of amplification and primer-binding bias. Taken together, our results provide a more unbiased perspective of abundances of microeukaryotes in global ecosystems.

Our eukaryotic census also provided a more detailed perspective on the distribution of eukaryotic groups across different latitudes (Figure 2D). Even at elevated taxonomic ranks, distinct patterns emerged across latitudes and ecosystem types. The marine environment exhibited both high prevalence and abundance of the Stramenopiles, Alveolata, and Rhizaria (SAR) group. Notably, at the arctic tundra around 90°, representatives from all three divisions were consistently present (prevalence = 1) and exhibited relative abundances of 5.85, 4.71, and 2.02 for Alveolata, Stramenopiles, and Rhizaria, respectively. Discobans were also found to be prevalent (prevalence = 0.93) but less abundant (relative abundance = 1.69) in close proximity to the North Pole. Prior studies using documentary sources, such as cell counts, showed the presence of alveolates (dinoflagellates), stramenopiles (diatoms), and discobans (euglenids) within the pan-Arctic region; however, rhizarians were not observed in this context (Poulin et al. 2011). Notably, the Euglenozoa subdivision was found with a very high mean abundance of 103.35 (SD=745.05) and a prevalence of 0.20 (SD=0.20) at 40° latitude, which agrees with previous observations that some groups of euglenids such as Diplonemida are the most abundant protists in marine environments (Singer et al. 2021). In freshwater ecosystems, Alveolates, particularly Ciliophora, displayed the highest mean prevalence (0.35, SD=0.36) alongside high relative abundance at 70° and 30° (relative abundance = 6.77, SD=21.40 and 4.40, SD=7.85, respectively), while groups such as Euglenozoa (Discoba) were also represented with mean prevalence = 0.20, SD=0.27 and high abundance at a latitude of 30° (relative abundance = 42.60, SD=1.39), with Discoba_X (primarily composed of heteroloboseans) exhibiting a mean abundance of 5.51 (SD=17.60).. Within soil ecosystems, Cercozoa (Rhizaria) persisted as the most abundant protist subdivision in soil with a mean abundance of 11.04 (SD=43.75), followed by Discoba_X (5.42, SD=12.28) and Chlorophyta_X (SD=5.41). This is in line with previous reports that showed that Cercozoans comprise a substantial proportion of microbial biomass in soil when evaluated using metatranscriptome sequencing data (Urich et al. 2008), often represented by highly divergent 18S rRNA gene sequences even among closely related lineages, as observed from isolates (Howe et al. 2011). Our results confirm that cercozoans represent a dominant eukaryotic group in soil environments with variable abundance as a function of latitude even within soil environments, and further expand the prevalence of this group through many newly identified lineages.

Our dataset of 18S rRNA gene sequences from thousands of samples presents a unique opportunity to estimate the taxonomic diversity of protists on a global scale. Rarefaction curves of the number of observed OTUs as a function of number of sequencing datasets showed that microbial eukaryote OTU richness did not reach a plateau for any of the ecosystem types, indicating that more extensive sampling efforts could yield a substantial number of novel OTUs (Figure 2E). Partitioning the samples among ecosystem types according to their GOLD ecosystem subtype showed that the highest eukaryotic diversity within marine systems was observed in epipelagic, oceanic, and intertidal zone subtypes (Figure 2F). The highest diversity in freshwater was found in lake and river subtypes, showing the highest diversity overall. Within soil type, peat and loam presented the highest diversity. This contrasts with a prior amplicon sequencing-based study which reported soil as the most diverse ecosystem for eukaryotic taxa (Singer et al. 2021). However, this previous research focused exclusively on topsoil and planktonic samples, while our study highlights that sediment samples represent important contributors to eukaryotic diversity in aquatic environments (Supplementary Figure 3). Freshwater ecosystems might provide nutrients for trophic groups to grow and in turn allow growth of heterotrophic organisms. These ecosystem dynamics align with the crucial role of nutrient cycling in supporting diverse trophic interactions in freshwater systems (Cotner and Biddanda 2002). These results indicate that nutrient availability and a stable ecosystem can in turn produce microbial communities as diverse as others in complex ecosystems such as the soil as freshwater bacterial communities often rival soil communities in phylogenetic diversity (Newton et al. 2011). Further, environmental stability and resource availability are key drivers of microbial community complexity across ecosystem types, including both aquatic and terrestrial environments (Delgado-Baquerizo et al. 2018).

Further, we were interested in patterns of the latitudinal gradient of eukaryotic diversity, as these patterns have been studied for other microbial groups such as marine bacteria (Fuhrman et al. 2008) and is a recurrent question in ecology (Mittelbach et al. 2007). We assessed the correlation between absolute latitude and OTU diversity within each eukaryotic subdivision across the three most prevalent ecosystem types in our samples. We were able to identify distinct patterns in the correlation of the 18S rRNA diversity with absolute latitude. Most taxonomic groups displayed positive coefficients in the quadratic term, indicating that diversity peaks at the Equator and the poles (Figure 2G). This finding contrasts with previous studies, done on observation data based on richness data of observations of benthic diatoms, suggesting that protist diversity shows weak or no correlation with latitudinal gradients, in contrast to plants and animals, which exhibit pronounced responses influenced by body size (Hillebrand and Azovsky 2001). Other investigations focused on specific taxa, such as Fungi, which exhibited a higher diversity at intermediate absolute latitudes (Bahram et al. 2018), in contrast to their metazoan counterparts. This was also evident in our analysis. Further, fungal lineages have been shown to have an increase in diversity near the equator, with tendencies towards reduced diversity in polar regions (Azovsky and Mazei 2013; Lara et al. 2016). While this trend was also visible in our data for some groups such as Apicomplexa in soils, we found opposing trends that presented higher diversity near the poles such as ciliates in soil and euglenids in freshwater.

### New Lineages of Eukaryotes

Our comprehensive census of protistan diversity has uncovered a substantial number of novel eukaryotic groups (2,002 OTUs at 85% sequence similarity) that are solely represented by metagenomic sequences, eluding classification within any known eukaryotic lineage. In order to resolve these unclassified lineages, we expanded our robust species tree utilized for taxonomic placement by incorporating these taxa.

The expanded tree preserved the majority of eukaryotic lineages as monophyletic, while also accommodating several orphan sequences interspersed among them (black branches, n=1,420) (Figure 3A). The presence of long orphan branches grouping within known eukaryotic phyla is noteworthy. A plausible explanation for these long branches is accelerated evolutionary rates, analogous to those documented in obligate intracellular bacteria (Ferla et al. 2013) and in eukaryotic parasites like microsporidians, which are known to exhibit highly divergent 18S rRNA gene sequences (Vossbrinck et al. 1987). While such branches can result from evolutionary processes, including rapid diversification or divergent molecular evolution, this pattern warrants further investigation using complementary approaches. For example, only through single cell transcriptome sequencing was it possible to identify a novel alveolate intracellular parasite closely related to perkinsids (Cooney et al. 2024). Cultivation of numerous eukaryotic lineages is notoriously challenging, leaving a considerable proportion of eukaryotic diversity unexplored on both genomic and phenotypic levels. In this study, we did not identify previously documented orphan OTUs, as our work mitigated primer binding bias. Indeed, the bias score exhibited significantly higher values (adjusted p-value = 0) in orphan sequences compared to reference sequences, irrespective of their origins from isolates or environmental sequencing datasets (Figure 3B).

**Figure 3.**
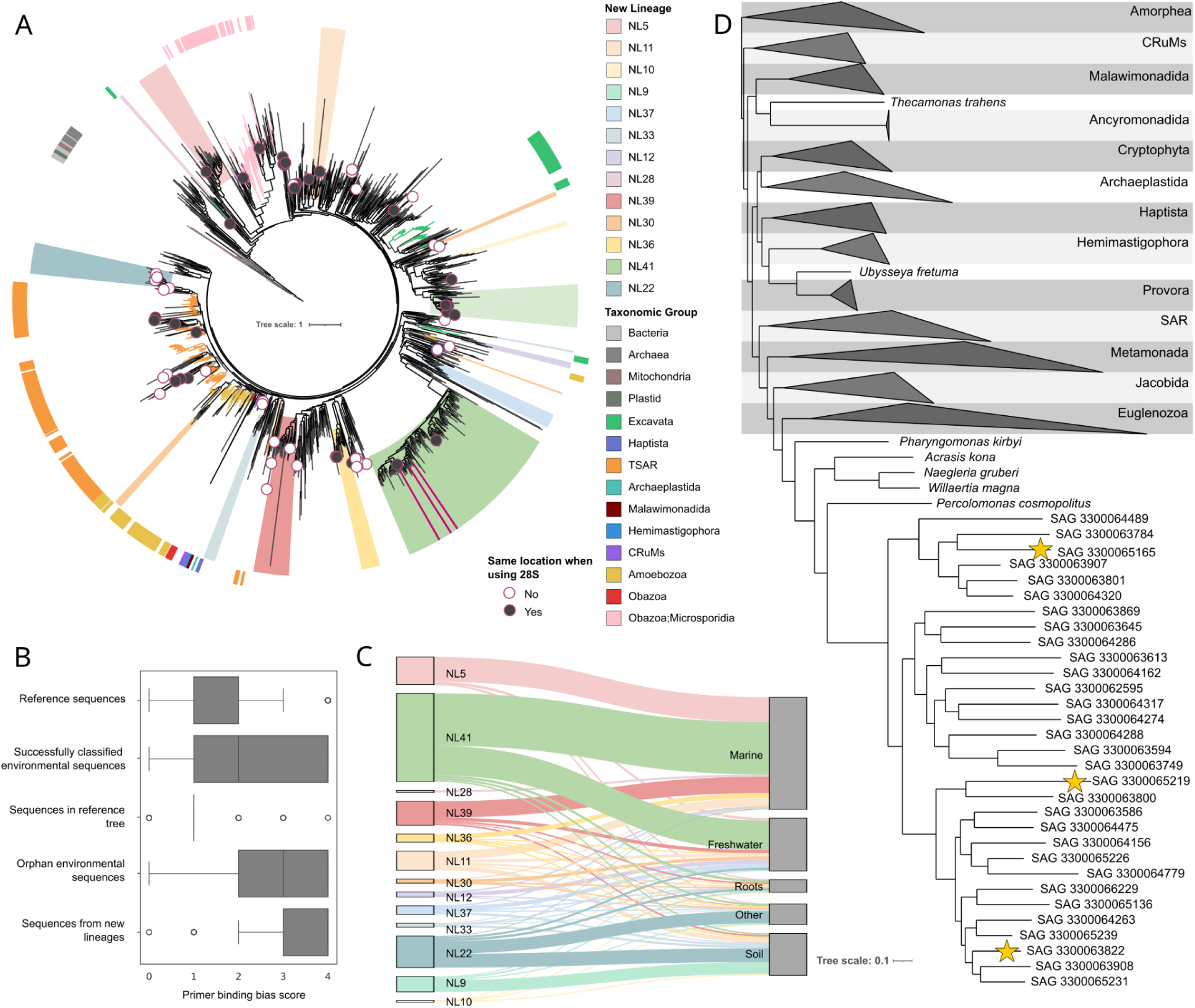
Distribution of new lineages of Eukaryotes. A) Maximum-likelihood tree derived from a concatenated alignment of 18S rRNA and 28S rRNA genes representing reference sequences (branches colored according to division-level taxonomy) and orphan 18S rRNA gene sequences (black branches). Clades likely indicative of major new lineages (NL) are highlighted with colored segments. Dark purple circles denote orphan contigs that encompass a 28S rRNA gene sequence. Filled circles indicate conserved placements maintained upon adding the LSU sequence to the multiple sequence alignment. Purple bars within NL41 represent SAG-derived 18S rRNA gene sequences. Clades belonging to Metazoa, Fungi and Streptophyta are collapsed. B) Distribution of primer binding bias scores across groups of 18S rRNA gene sequences aggregated from our combined database. All comparisons exhibited an adjusted p-value = 0 in paired Mann-Whitney tests. Boxes show interquartile ranges, lines indicate medians, whiskers represent data spread, and circles denote outliers. C) Diagram illustrating the GOLD Ecosystem type category assigned to the sequencing samples containing sequences from the newly identified lineages. D) Maximum likelihood tree based on a concatenated alignment of 341 protein sequences. Stars denote SAGs from which 18S rRNA gene sequences were extracted, branching within NL41 in the phylogenetic tree represented in A.

Notably, the majority of environmental orphan sequences (n=1,420) branched deeply in the eukaryotic tree or in proximity to discoban sequences bridging excavates and the eukaryotic root. We identified 13 clades entirely composed of orphan OTU representatives (466 in total), likely indicative of major, undescribed new lineages (NLs, colored ranges). Given that 28S rRNA gene sequences were also detected within some metagenomic contigs containing orphan 18S OTU representatives, we examined whether the inclusion of additional phylogenetic information through adding the 28S rRNA gene sequences would impact their classification and placement within the reference tree. In our results the placement remained consistent for sequences in NL11, NL9, NL41, and several orphans within TSAR (filled red circles). New lineages 5 and 11 branched outside of all eukaryotic sequences from the reference tree, suggesting they may represent early branching eukaryotes. In contrast, several of these new clades were affiliated with Excavata, specifically the Discoba division, such as NL9. Various clades branched as sister groups to sequences from the discoban phylum Heterolobosea: NL37, NL33, NL12, NL28, NL39, NL30, NL36, and NL41. The most phylogenetically diverse group within these new lineages, NL41, formed a substantial clade that may represent a new discoban lineage closely related to Heterolobosea.

We further validated NL41 as a distinct lineage using an independent dataset of single-amplified genomes, derived from cells isolated from anoxic marine sediments in Maine. Phylogenomic analyses using a set of 341 conserved eukaryotic proteins indicated that the SAGs formed a novel clade, sister to *Percolomonas cosmopolitus* (Figure 3D). Within the novel SAG clade, we found three SAGs containing 18S rRNA gene sequences, all of which group within the NL41 clade in the 18S tree (Figure 3A,D). This phylogenetic placement, together with the observation that the best BLAST hit from the nucleotide database corresponding to these sequences was the 18S rRNA sequence of *Creneis carolina* (mean identity %=78.3), implies that NL41 belongs to Heterolobosea, potentially related to Tetramitia.

We traced the ecosystem types from which sequencing datasets belonging to the new lineages were obtained and observed that marine ecosystems contributed the largest proportion, followed by freshwater and soil (Figure 3C). Upon assessing the environmental sources of these datasets, we found that wetlands from both marine and freshwater ecosystems were the most frequently represented ecosystem subtypes (Supplementary Figure 4; Supplementary Table 4). Wetlands harbor diverse microbial communities, including the newly proposed bacterial phylum Candidatus Cosmopoliota (Zhang et al. 2023), which was identified through metagenome-assembled genomes (MAGs) recovered from this ecosystem.

We used an example lineage for which there was limited taxonomic confidence to demonstrate that more than just the 18S rRNA gene could be extracted for these novel lineages from the metagenomes. We chose the novel lineage NL5 because of its location in the tree among discoban clades and other groups composed of unclassified sequences and better classify it in the eukaryotic tree using recovered coding genes, and additionally applied the pipeline to NL11, which appeared to be exceptionally novel. Thus we applied the tax-aliquots clustering approach (described in the Methods) to identify functionally annotated genes that clustered with the non-18S rRNA gene portions of contigs containing the 18S rRNA gene established through our discovery clustering pipeline (Krinos et al. 2024). This approach clusters sequences based on their *k*-mer frequencies to identify sequences from similar taxonomic lineages that cannot be aligned to existing reference sequences. We retained clustered contigs associated with protein family annotations (Pfam database) with e-values less than 10^-5^ via HMM search, thus stringently excluding clustered sequences lacking robust protein annotations. The presence of functional genes within nucleotide sequences which clustered with the 18S rRNA gene-containing contigs provides additional evidence that the 18S rRNA genes are valid.

Among sequences exclusively clustering with NL5, we identified 35 clustered contigs containing a valid Pfam annotation using the hierarchical clustering approach. These Pfam annotation-containing contigs included multiple membrane proteins (amino acid permease and ABC membrane transporter proteins), eukaryotic flagellar proteins, DNA replication proteins, glutamate synthase, and cytochrome p450 (Supplementary Table 5). We aligned these functionally-annotated proteins with proteins from the nr database from NCBI (Pruitt et al. 2006) to identify functionally-annotated proteins with similar taxonomic alignment position to NL5. Of 45 unique Pfams annotated across 54 distinct proteins identified in nucleotide-clustered contigs, 24 of 54 (44%) were taxonomically annotated as eukaryotes and 9 (38%) of those eukaryotes had a maximum percentage identity of less than 80% assigned to the best hit to the NR database. 21 protein sequences (39%) did not have any BLAST hit to the NR database at all. This result aligns with the novel diversity identified via the 18S rRNA gene sequences and provides putative functional proteins that appear to have derived from the new lineages. We applied the same pipeline to NL11 and focused only on sequences in hierarchical clusters containing only NL11 rRNA gene-containing contigs as well as contigs which did not contain a new lineage rRNA gene nor a ribosomal functional annotation. Only retaining hits with e-value <10^-5^, we identified 500 proteins containing 357 Pfam annotations among contigs from samples in the IMG/M database (Supplementary Table 6). These taxonomic annotations included ABC membrane transporter proteins, phosphomethylpyrimidine kinase, acetyltransferases, and fatty acid hydroxylases. We found through initial taxonomy screening that many of these sequences had uncertain taxonomy, so we annotated the sequences with a secondary database and found that 86 (17.2%) had all top 10 hits eukaryotic, 66 (13.2%) had all top 10 hits bacterial, 23 (4.6%) had no clear hits to the taxonomy database, and 325 (65%) had both eukaryotic and bacterial hits in the top 10. Hence, most of the protein sequences that clustered with masked rRNA gene-containing contigs were of ambiguous taxonomic origin.

The position of NL11 in the 18S-28S rRNA gene tree among orphan sequences and separated from known eukaryotic supergroups could point to a distant phylogenetic relationship with most eukaryotic lineages, even within another domain of life. Our tree-distance analysis indicates NL11 sits closer to eukaryotes than to bacteria, archaea or organelles, consistent with a highly divergent eukaryotic lineage; given the mixed taxonomic affinities of its associated proteins, we treat this placement as provisional pending genomic confirmation (Supplementary Figure 6). These divergent rDNA sequences could reflect a particular evolutionary history producing high substitution rates as found in rhizarian lineages (Burki and Pawlowski 2006) as well as highly modified genomes associated to a specific lifestyle such as the reduced genomes of intracellular symbionts such as the parasitic microsporidia (Vossbrinck et al. 1987).

Previous amplicon-based studies of eukaryotic diversity in environmental samples have indicated that many protist groups contain representatives that remain virtually unknown (de Vargas et al. 2015). The classification of these new clades poses challenges, often limited to specific clades (Reboul et al. 2019; Fermani et al. 2021), primarily due to the conventional short lengths of amplicon libraries, which cover only selected regions of the 18S rRNA gene (Hugerth et al. 2014). To overcome some of these limitations, we recovered 18S rRNA genes (>1,200 bp) from assembled contigs and, if co-located, also the 28S rRNA genes of novel eukaryotic lineages, improving phylogenetic resolution. Our findings imply the presence of at least 13 major clades of eukaryotes lacking both genomic representation and closely related sequences in reference databases, representing taxonomic blind spots. We also show how these groups can be placed in the tree of eukaryotes with confidence using single-cell sequencing methods. The detailed information on sample origins provided in this study, predominantly wetlands and sediment systems, will facilitate future targeted sampling efforts aimed at further exploring this unexpected novelty within the eukaryotic tree of life.

We mapped approximately 166,000 short-read V4 18S rRNA gene amplicon sequences against our database of ∼13,000 metagenomic OTU representative sequences. Across the merged EukBank and MetaPR2 V4 datasets, 30.94% of our metagenomic OTUs matched an amplicon sequence and 50.33% of short amplicon OTUs matched our metagenomic OTUs (Supplementary Figure 5; Supplementary Dataset); part of this gap reflects habitat differences between the amplicon datasets and IMG/M (detailed below). This disparity was particularly pronounced for novel lineages. For instance, only 21 OTUs from 5 orphan major clades (NLs) aligned with short-read sequences, which were found in low abundance in amplicon sequencing datasets with a median abundance of 52 (mean ASV abundance=1120.97, SD=4623.22).Thus, the novel lineages we identified from metagenome assemblies have been largely missed by the amplicon-based studies cataloged in the EukBank dataset and MetaPR2 database (Berney et al. 2023; Vaulot, et al. 2022). Further supporting this conclusion, we observed a small but significant negative Spearman correlation (rho = −0.19, p = < 0.001) between the weighted environmental abundance of OTUs and their calculated mean primer-binding bias score. This limited shared genetic diversity also arises from comparing datasets from different sources: The EukBank and MetaPr2 datasets agglomerate samples from specific environmental types, such as marine, freshwater and soil. In contrast, the IMG/M database, from which most environmental sequences were derived, also includes host-associated samples such as digestive systems and built environments such as anaerobic digesters. For instance 3,114 of the 12,398 samples containing protist 18S sequences from the IMG/M database have a GOLD environmental type annotation different from Marine, Freshwater, and Soil. These results show where amplicon-only surveys fall short and why metagenome assembly is needed to explore eukaryotic diversity, especially for the discovery and phylogenetic placement of novel groups.

### Prediction of Associations Between Microbes Across the Globe

Microbial eukaryotes are documented to coexist with other microbes, some serving as hosts for bacterial symbionts, which may function as either ecto- or endosymbionts. These relationships can provide nutrient benefits but may also confer defense mechanisms and/or environmental adaptability (Gast et al. 2009; Husnik et al. 2021). Our abundance correlation results predict associations among unclassified eukaryotes and bacteria annotated as Verrucomicrobia, ABY1, and unclassified bacteria in marine ecosystems (n=7, 4, and 3, respectively; Figure 4A). OTUs categorized within various families of the Amoebozoa supergroup demonstrated a relatively high incidence of positive correlations with prokaryotic OTUs identified as Vicinamibacteraceae (n=15), Ardenticatenales (n=25), and Bacteroidiota (n=13) in freshwater ecosystems (Figure 4B), as well as Rickettsiales (n=5) and Microgenomata (n=4) in soil environments (Figure 4C). Several predicted associations likely represent genuine symbioses. This is exemplified by interactions involving Rickettsiales bacteria (identified via BLAST: >80% identity, >700 bp alignment length), a lineage well-documented as endosymbionts of amoebozoans, algae, and ciliates (Schulz et al. 2025; Schulz et al. 2016; Lanzoni et al. 2019). Notably, cell-resolved studies have directly recovered genomes of bacteria physically co-associated with uncultivated marine predators, supporting the plausibility of stable protist-bacteria partnerships. (Needham et al. 2022; Wittmers et al. 2025). The congruence between these inferred interactions and established ecological relationships supports the biological relevance of our predictions. Other associations likely reveal environmental or feeding preferences, as suggested by prior studies (Oliverio et al. 2020).

**Figure 4.**
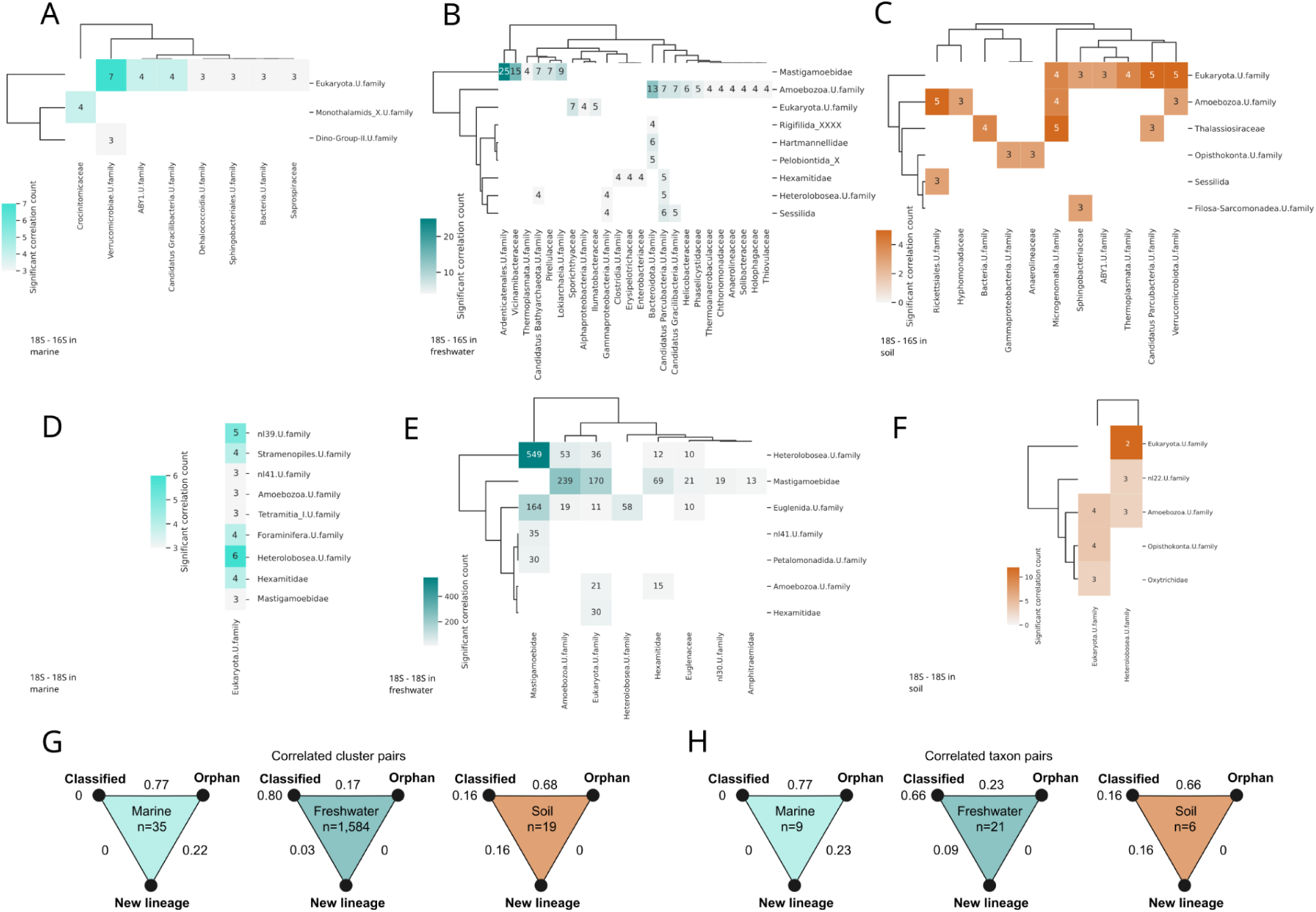
Association between abundances of protists and prokaryotes across major ecosystem types. Heatmaps illustrating the count of significant positive correlations between 18S and 16S rRNA gene sequences (A, B, and C) as well as between 18S and 16S rRNA sequences (D, E, and F) aggregated at the family level. Values are represented for family pairs exhibiting a minimum of three significant (adjusted p-value < 0.05) correlations. G. Proportions of the total number of correlated pairs (n) between each cluster type according to their taxonomic classification (Classified, Orphan, New Lineage) for each ecosystem type regarding abundance correlations among eukaryotic sequences. H. Proportion of the total number of taxa manifesting correlated pairs between each cluster type across each ecosystem type.

Correlations among various eukaryotic lineages within marine environments also revealed possible associations between unclassified eukaryotes at the supergroup level and unclassified sequences associated with groups such as Heterolobosea (n=6), NL39 (n=5), and NL41 (n=3; Figure 4D). Freshwater ecosystems exhibited the highest number of correlations overall (n=1,584), with sequences from Mastigamoeba and Heterolobosea representing 549 pairs (Figure 4E). The anaerobic growth preferences of many amoebae within these groups, particularly members of the Psalteriomonadidae family, coinciding with the anaerobic lifestyle of some heteroloboseans (Pánek et al. 2012) may account for this finding. We also identified 19 eukaryote-eukaryote co-occurrences across the soil samples, primarily involving unclassified groups of Heterolobosea and eukaryotes unclassified at the supergroup level (Figure 4F). Some members of the new lineages identified in this work were found correlated with other eukaryotes, such as NL39 and NL41 correlated with unclassified eukaryotes in marine ecosystems, NL30 and NL41 with Mastigamoebidae in freshwater, and NL22 with heteroloboseans in soil. These associations could represent host-parasite interactions considering the highly divergent sequences of these new lineages, which is often a signature of an intracellular lifestyle (Vossbrinck et al. 1987). Rhizarian endobiotic parasites are known to infect other protists, such as *Woronina pythii* (Cercozoa) which infects several species of *Pythium* (Oomycota) (Goldie-Smith, 1956). Similarly, predatory protists are known to prey upon both bacteria and other eukaryotes, and certain species of amoebozoans and heteroloboseans exhibit species-specific feeding habits (Berlinches de Gea et al. 2024). Our findings could facilitate the identification of more eukaryotic species-specific predatory relationships between members of these recognized groups and the newly identified eukaryotic lineages.

The proportion of correlated OTU pairs, whether classified to an existing supergroup, orphan, or assigned to a new lineage, varied across ecosystem types (Figure 4G). In marine and soil ecosystems, a predominance of associations occurred between classified and orphan OTUs (0.77 and 0.68, respectively), while in freshwater systems, the majority of correlated pairs comprised two successfully classified OTU representatives (0.80). This trend became even more evident when comparing the proportions of taxon pairs with correlated pairs (Figure 4H), noting a reduced ratio of taxon pairs classified at the supergroup level in freshwater (0.66). Likewise, we found a reduced proportion of associations between orphan taxa and taxa classified at the supergroup level in soil (0.66). Within marine ecosystems, the relationships between orphan OTUs and those classified as new lineages represented the second-largest proportion (0.22 and 0.23 for correlated OTU pair count and taxa with correlated pairs, respectively). Established routine sampling methods, often optimized for water quality monitoring and pathogen detection, seem to have led to an increased effort but also bias in the exploration of protist taxa in freshwater ecosystems (Dodds, 2005). Soil has been characterized as the ecosystem type with the highest novelty regarding documented 18S rRNA gene sequences and marine the ecosystem type with even less novelty than freshwater ecosystems (Singer et al. 2021). Thus, the more frequently recovered predicted associations between known taxa in freshwater can likely be attributed to the bulk of previous work, alongside the fact that the taxa exhibiting the most associations (amoebozoans and discobans) are found to be phylogenetically diverse and may establish relationships with other eukaryotes. Conversely, the higher proportion of associations involving known and orphaned taxa that were found in this study underscores the significance of associations formed by understudied protist groups within marine and soil ecosystems.

## Conclusion

Our comprehensive, metagenome-based survey of eukaryotic diversity substantially expands the known breadth of protistan lineages and reveals previously unseen branches across the eukaryotic tree. By leveraging assembled metagenomes to circumvent the inherent biases of amplicon-based approaches, we captured novel lineages belonging to taxonomic blind spots that often lack cultured representatives and inhabit a broad range of global environments. The identified protists include largely divergent groups within Excavata and previously uncharacterized taxa branching deeply in the eukaryotic tree. We further uncovered likely associations between understudied protists and bacteria, hinting at intricate symbiotic or predator-prey relationships that shape community structure. These results give targeted cultivation, genomic, and ecological studies a concrete set of lineages and sampling locations to pursue, and they expand the known branches of the eukaryotic tree that current databases miss.

## Methods

### rRNA Sequence Search

We compiled assembled contig files corresponding to 32,520 public datasets derived from metagenome, metatranscriptome, and single-cell sort sequencing projects marked as both public and unrestricted within the Integrated Microbial Genome/Metagenome (IMGM) database. The cmsearch program from the Infernal v1.1.5 (Nawrocki and Eddy 2013) package was utilized to search the contig files for sequence matches to each of the following small subunit rRNA (SSU rRNA) models from the Rfam database (Kalvari et al. 2018): eukaryotic 18S rRNA (RF01960), microsporidial 18S rRNA gene (RF02542), archaeal 16S rRNA gene (RF01959), and bacterial 16S rRNA gene (RF00177). Employing the --anytrunc option in cmsearch allowed for the inclusion of long SSU rRNA sequences in our analysis, enabling us to parse the output for contigs displaying sequential and multiple non-overlapping alignments to the covariance models (Supplementary Methods). Furthermore, long subunit rRNA (LSU rRNA) sequences were screened using the eukaryotic 28S rRNA gene (RF02543), archaeal 23S rRNA gene (RF02540), and bacterial 23S rRNA gene (RF02541) covariance models from Rfam, utilizing cmsearch with default parameters.

### Preliminary Annotation

We searched the retrieved SSU rRNA sequences against a database that contained all SSU rRNA sequences from both Silva (Quast et al. 2013) and PR2 (Guillou et al. 2013) databases using the BLAST+ v2.13.0+ (Camacho et al. 2009) program with an evalue cutoff of 0.005 and considering hits with sequence identity % > 70 to account for potential divergent sequences. Sequences with no hits to the database were added to a reference SSU rRNA tree to confirm their phylogenetic affiliation at the domain level. The reference tree was built as follows: SSU rRNA from the Silva and PR2 databases were clustered at 75% identity to group higher groups and further reduce redundancy using Usearch v11.0 (Edgar 2010) and the resulting centroids were aligned with Mafft v7.525 (Katoh and Standley 2013) using the Linsi algorithm. Tree building was done with IQTree v1.6 (Nguyen et al. 2015) with default parameters and model GTR+F+R10. Sequences that branched within a cluster that belonged to a different domain were considered misannotations and subsequently removed from the alignment and the tree then rebuilt with the filtered alignment. Sequences confirmed by tree building to belong to the domain Eukarya were added to the 18S rRNA gene sequence database together with those that had a best hit with 18S rRNA gene sequences in Silva and PR2. We retained sequences with a length >= 800 bp, as this threshold represents approximately half of the SSU rRNA gene and is sufficient to capture the conserved regions essential for robust phylogenetic analyses. This search and filtering protocol yielded 157,956 18S rRNA gene sequences derived from 18,445 distinct datasets.

### rRNA SSU Sequence Clustering

We assembled a database inclusive of 18S rRNA gene sequences from the Silva database (Quast et al. 2013), the PR2 database (Guillou et al. 2013), sequences referenced in the description of the Provora supergroup (Tikhonenkov et al. 2022), long amplicon sequences from a recent study (Jamy et al. 2022), sequences extracted from eukaryotic genomes housed in GenBank (Sayers et al. 2022) represented within the EukProt database (Richter et al. 2022), as well as environmental sequences sourced from IMG/M in February 2022 (Chen et al. 2023). Two-step clustering was conducted using the Usearch program with the -cluster_fast option, initially targeting 97% identity and subsequently at 85% identity to account for the high variability of 18S rRNA gene sequences in related taxa. Following the removal of clusters containing only a single member, we obtained 13,238 sequence clusters (OTUs), wherein the largest cluster comprised 18,608 sequences.

Concurrently, SSU rRNA sequences extracted from the IMG/M database, those recognized as having a 16S rRNA gene as their best hit during BLAST searches, were clustered at 97% identity and subsequently at 85% together with the 16S rRNA gene sequences from Silva and PR2 databases. Clusters with only a single member were excluded from analysis. Taxonomic annotations for each sequence were inferred from the cluster representative’s best hit and reformatted to be compatible with the NCBI Taxonomy structure. This procedure produced a total of 839,222 16S rRNA gene sequences divided among 23,233 clusters, representing Bacteria, Mitochondria, Archaea, and Plastids (Supplementary Figure 7).

### Primer Binding Analysis

The software PrimerProspector v1.0.1 (Walters et al. 2011) was employed to search for four different primer pairs across each 18S rRNA gene sequence from the non-singleton 85% identity clusters. Specifically, we employed fwf/rvr (Nolte et al., 2010) for the V3–V4 region (yeast positions 366–586); 616of/1132r (Hugerth et al., 2014) for the V4–V5 region (positions 616–1132); and 1380F/1510R (Amaral-Zettler et al., 2009) for the V9 region (positions 1625–1770). Additionally, we included the Fwd1f/Rev3r pair (Stoeck et al., 2010), which targets the V4 region (positions 564–981). This latter represents the most widely used primer pair in global biodiversity initiatives. For instance, it accounts for approximately 80% of the currently available sequence data deposited in EukBank (Berney et al., 2023). Sequences with an overall weighted score exceeding 1 for any forward or reverse primer were classified as undetected by respective primer pairs. We aggregated the weighted score of all four primer pairs to calculate the binding bias score and subsequently summarized the binding bias score at the cluster level, determining the mean bias score of all sequences within each cluster. We compared the distribution of primer binding bias scores across different dataset groups: reference sequences (originating from Silva, PR2, Provora, Genbank, and long amplicons), successfully classified environmental sequences, orphan environmental sequences (lacking classification at the supergroup level), and orphan sequences representing potential new lineages. To assess significance, we performed a Kruskal-Wallis test, followed by paired two-tailed Mann-Whitney tests across pairs, adjusting p-values via the Benjamini-Hochberg procedure using the scipy.stats module v1.16 (Virtanen et al. 2020). A p-value < 0.05 was deemed statistically significant. Resulting effect sizes, p-values and adjusted p-values are available in Supplementary Table 7.

### 18S and 28S rRNA Gene Tree Construction

We selected a subset of 1,247 cluster representative sequences based on specific inclusion criteria: 1) both the 18S and 28S rRNA gene sequences could be extracted from the same contig; 2) 18S sequences exhibited lengths of >= 1,200 bp; 3) 28S sequences displayed lengths of >= 2,800 bp; and 4) centroids were sourced from environmental samples from IMG or from long amplicon sequencing (Jamy et al. 2022). Employing the cmalign program and the RF01960 model, we aligned 18S rRNA gene sequences, and utilized the RF02543 model to align 28S rRNA gene sequences, employing the --match-only option to restrict alignment sites to those represented within the model. We concatenated sequences from both rRNA genes utilizing AMAS v1.0 (Borowiec 2016) and constructed a tree through IQTree applying the GTR+F+R10 model.

To enhance the tree, we removed terminal branches exhibiting lengths > 0.5. We then recovered the original versions of the remaining sequences and separately aligned the 18S and 28S rRNA gene sequences using mafft-linsi software. Subsequently, we concatenated the alignments of both genes and employed the trimAl v1.4 program (Capella-Gutiérrez et al. 2009) with the -gt 0.1 option to exclude sites with significant indel content. Afterwards, we used the IQTree program with the GTR+F+R10 model along with 1,000 ultrafast bootstrap replicates to construct a phylogenetic tree. The resultant tree underwent additional manual inspection in which sequences branching within a cluster dominated by sequences from differing supergroups were excluded from the concatenated alignment. The aforementioned removal rogue taxa has the trade-off of masking true diversity that could result of misclassification of some groups, but improves tree stability for downstream analyses (Aberer et al. 2012). The final tree included 577 sequences.

### Phylogenetic Placement Based Annotation

We selected an outgroup comprising 38 bacterial, archaeal, and organellar genomes from GenBank (Supplementary Table 8), aligning the respective SSU and LSU rRNA gene sequences with the RF01960 and RF02543 covariance models utilizing the cmalign program with the –match-only and --ifile options. A phylogenetic tree was developed using sequences from the eukaryotic 18S and 28S tree alongside outgroup SSU and LSU rRNA sequences, constructed by IQTree employing the GTR+F+R10 model and incorporating 1,000 ultrafast bootstrap replicates. All cluster representatives of 18S rRNA gene sequences were aligned with the RF01960 covariance model using the --match-only option, subsequently appending a tail of gaps (-) to each sequence to account for the absence of the corresponding 28S rRNA gene sequence. We utilized the EPA-NG v0.3.8 software (Barbera et al. 2019) to place aligned centroid sequences into the reference tree with the inclusion of the added outgroup, assigning taxonomy to the placed sequences based on the reference tree taxonomy via the Gappa v0.9.0 software (Czech et al. 2020). We implemented a hierarchical matching algorithm to harmonize

Silva annotations with the PR2 taxonomic framework. This adopted framework designates clades of unconfirmed monophyly using an “_X” suffix (Guillou et al. 2013). The algorithm first queried the PR2 database for an exact match to each Silva annotation at its lowest taxonomic rank. If a match was identified, the complete higher-level PR2 classification was inherited. If no match was found at the lowest rank, the algorithm iteratively ascended the Silva taxonomic hierarchy, searching at each progressively higher rank until a corresponding entry was found in PR2, at which point that classification was assigned. To account for truncated taxonomic assignments typical of phylogenetic placement, we implemented a systematic nomenclature for unresolved ranks. OTUs that could not be classified at a specific level were assigned a suffix consisting of “.U.” (Unassigned) followed by the rank name. For example, an OTU phylogenetically placed within Euglenozoa but lacking class-level resolution was designated “Euglenozoa.U.class”; this convention was applied recursively to all subsequent downstream ranks (e.g., “Euglenozoa.U.order”).

### Chimera Detection and Filtering

To discard potential chimeric sequences produced by metagenome sequence assembly, we subjected our cluster representative sequences of the 13,238 18S OTUs and 23,233 16S OTUs separately to a two-step chimera detection procedure. We used vsearch v2.29 (Rognes et al. 2016) with both the de novo chimera detection algorithm (--uchime_denovo) and with the reference-based algorithm (--uchime_ref) (Edgar et al. 2011). As references for the latter, we used 241,792 18S rRNA reference sequences, from Silva, PR2 and Provora, and 459,898 16S rRNA reference sequences from Silva and PR2 for the 18S and 16S cluster representative datasets, respectively. This procedure identified 812 18S and 2,462 16S chimeric sequences, of which only 9 were part of NLs, all of which were excluded from downstream analyses.

### 18S rRNA Insertion Distribution Analysis

Using the per-cluster 18S rRNA representative sequence insert information produced by the cmalign program with the RF01960 covariance model and the --ifile option, we recovered only those inserts with length >= 10 bp. Then we obtained and compared the distribution of both the number of inserts and length of inserts for each eukaryotic division. To assess significance, we performed a Kruskal-Wallis test, followed by paired two-tailed Mann-Whitney tests across pairs, adjusting p-values via the Benjamini-Hochberg procedure using the scipy.stats module v1.16 (Virtanen et al. 2020). A p-value < 0.05 was considered statistically significant. Resulting effect sizes, p-values and adjusted p-values are available in Supplementary Table 9 and Supplementary Table 10 for differences in insertion number and length, respectively.

### Global Distribution Analysis

We extracted longitude and latitude data from the IMG/M database metadata and visualized it through the geopandas module for Python3. Latitude data was digitized into a discrete scale with a step size of 10 degrees. Distribution was visualized using the geopandas module (Jordahl et al. 2022) for Python3. Abundance was defined as the average of the normalized mean sequencing depth across all contigs pertaining to each taxonomic group, while prevalence was defined as the proportion of samples from comparable environments and latitudes in which each taxonomic group was identified. For the evaluation of eukaryotic family distribution, we focused exclusively on those groups attaining prevalence and abundance at or above the 80th percentile to emphasize those groups that were highly abundant and prevalent.

### Sequence Read Depth

We retrieved data on per-contig average read depths across metagenomic datasets whenever available from IMG/M, employing this information to estimate the sample-wide average read depth via the following expression:

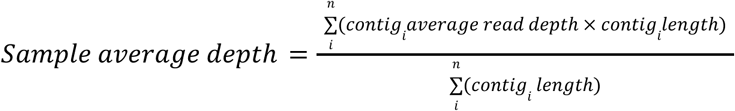

The sample average depth was subsequently employed to compute the normalized average read depth for every contig harboring an SSU rRNA gene in the dataset.

### Ecosystem Diversity Analysis

For this analysis, we exclusively considered samples from the three most prevalent GOLD ecosystem types represented in our dataset: Marine, Freshwater, and Soil (as defined by GOLD ecosystem type metadata; ‘Soil’ includes peatlands/wetlands and other soil-associated terrestrial subtypes). Given that the metrics utilized for analyzing diversity rely on count data, we simulated read counts based on the normalized average read depth data, employing the expression:

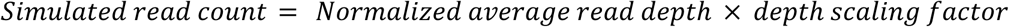

With a depth scaling factor of 100 to take into account sequences that were represented at least at 1% of the sample average depth, read counts were rounded to the nearest integer. Contigs with a simulated read count of 0 were subsequently removed. To account for sequencing disparity, we executed a random rarefaction employing the rrarefy function from R version (R Core Team, 2013) utilizing the vegan package (Oksanen et al. 2013) to align read counts with that of the smallest sample.

To visualize the 18S rRNA gene OTU sampling effort, we employed a per-sample basis rarefaction curve for each distinct ecosystem type, estimating the mean observed OTUs per new sample across 10,000 randomizations. We then fitted our accumulation curves to various linear and nonlinear models, as outlined below:

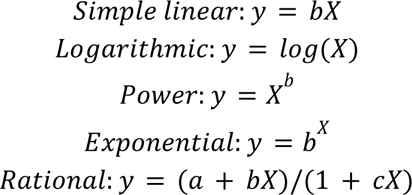

We used the Python ols function from the statsmodels module (Seabold and Perktold 2010) and the curve_fit function from the scipy module v1.16 (Virtanen et al. 2020) for model fitting. The best-fitting model for the accumulation curve was determined using the Akaike Information Criterion (AIC). The rarefaction curves for the Freshwater and Marine ecosystem types were comparable, exhibiting overlaps. We fitted the rarefied count data to five distinct models, inclusive of an asymptotic rational model. Across all assessments, the power function constituted the best fit, confirming a non-asymptotic relationship (Supplementary Table 11).

Subsequently, we utilized the Python3 ecopy module (https://github.com/Auerilas/ecopy) to quantitate Shannon H, Simpson, and Inverse Simpson diversity indices on the rarefied OTU table. Second-order polynomial linear regressions were applied to model the relationship between diversity (measured by the Simpson index) and absolute latitude. For this analysis, we segmented the OTU table for each taxon across varying taxonomic levels: supergroup, division, and subdivision, limiting our focus to taxonomic groups present in a minimum of 10 samples. To account for data partitioning, we adjusted p-values for polynomial fits using the Benjamini-Hochberg procedure.

### Search for Potential New Lineages

We constructed a phylogenetic tree utilizing OTU representative sequences classified under the eukaryotic domain and categorized unknown supergroups. These sequences were combined with data from the reference tree employed for phylogenetic placement analysis. Using concatenated alignments for the 18S and 28S rRNA genes based on the RF01960 and RF02543 covariance models, we constructed the tree employing the IQTree program with the GTR+F+R10 model and 1,000 ultrafast bootstrap replicates. Clades comprising at least five leaves classified as eukaryotic with unknown supergroup status and belonging to clusters containing only metagenomic sequences, along with maximum bootstrap support values, were defined as candidates for potential new lineages. The environmental origins of samples harboring sequences from these new lineages were visualized employing the plotting capabilities of the plotly v5.12 Python3 module (https://github.com/plotly/plotly.py). To compare the binding bias distribution among categories of sequences defined by their affiliation to new lineages, source type and presence in the 18S-28S reference tree, we grouped the average binding bias scores of 18S rRNA gene OTUs in the following categories: Reference sequences (derived from Silva and PR2), Successfully classified environmental sequences, Sequences in reference tree (environmental sequences used to build the reference tree), Orphan environmental sequences, and Sequences from new lineages. Then we performed a Kruskal-Wallis test, followed by paired two-tailed Mann-Whitney tests across taxonomic group pairs, adjusting p-values via the Benjamini-Hochberg procedure using the scipy.stats module v1.16 (Virtanen et al. 2020). A p-value < 0.05 was deemed statistically significant. Resulting effect sizes, p-values and adjusted p-values are available in Supplementary Table 12. Similarly, we compared tree distance distributions between NL11 and other taxonomic groups to clarify its association between different domains of life. We grouped distance values according to taxonomic groups (i.e. Archaea, Bacteria, Mitochondria, Plastids, and each eukaryotic supergroup) and performed the same *ad hoc* and *post hoc* tests as mentioned above. Effect sizes and p-values for the branch length analysis are available in Supplementary Table 13.

### Phylogenomic Analysis

To further validate the taxonomic affiliation of the potential new lineages, we used an independent dataset of single-amplified genomes derived from cells isolated from anoxic marine sediments in Maine. We performed a phylogenetic analysis expecting a robust classification based on several marker genes. Initial filtering included the removal of SAGs with fewer than 100 putatively annotated eukaryotic proteins. The peptides from the remaining SAGs were incorporated into broadly conserved gene families using PhyloToL v4.3 (Cerón-Romero et al. 2019), identifying 345 gene families well represented among the remaining SAGs. Subsequently, we excluded SAGs with low representation within this pool of gene families (i.e., present in <80 gene families), thereafter generating single-gene alignments, conducting homology assessments, and selecting orthologs with PhyloToL. For each alignment, columns with more than 90% missing data were removed with trimAl (Capella-Gutiérrez et al. 2009). Following alignment processing, we concatenated the 341 single-gene alignments, utilizing IQTree2 v2.2 (Minh et al. 2019) for phylogenomic reconstruction, applying the LG+C60+G+F model under the PMSF approximation with 100 bootstrap iterations.

### Tax-aliquots Clustering Method

The tax-aliquots clustering method (Krinos et al. 2024) was employed to revisit the original metagenomes, recovering contigs akin to the non-18S rRNA gene portions of the 18S rRNA gene-containing contigs used for identifying new lineages. We utilized mmseqs2 (version 13.45111) (Steinegger and Söding 2017) to precluster all contigs from samples from which a new lineage was identified, establishing a threshold coverage value of 0.01 and a percent sequence identity of 0.7 for NL5 and a percentage sequence identity of 0.97 for NL11 due the higher number of matches, utilizing coverage mode 1; we then performed a secondary hierarchical clustering of the preclustered sequences using a cutoff of 0.3 as described in Krinos et al. (2024). All contigs within clusters containing at least one sequence inclusive of an 18S rRNA gene-containing contig were retained, thus minimizing the total input size of sequence content while limiting sequence representatives to a manageable number by restricting the search to contigs already exhibiting some degrees of similarity to the rRNA gene-containing contigs. Subsequent to obtaining this original clustered assembly, we differentiated sequences associated with the original 18S rRNA gene-containing contigs from those that did not incorporate these genes. With the coordinates of the rRNA gene within the 18S rRNA gene-containing contig, we masked the 18S rRNA gene sequences, recombining the files as input for tax-aliquots. The reference content consisted of the masked 18S rRNA gene-containing contigs, while query content comprised other contigs remaining subsequent to the initial mmseqs2 filtering (employing SeqIO from Biopython for sequence masking; (Cock et al. 2009). We subsequently clustered this sequence content utilizing mmseqs2 v13.45111 (Steinegger and Söding 2017) with a coverage of 0.2 and percent sequence identity of 0.2, retaining clusters that encompassed at least one of the masked rRNA-gene containing contigs and at least one additional contig that did not contain an 18S rRNA gene. The resulting assembly was subject to further clustering with sequence information parsed into k-mers of length 3.

### Short Amplicon Sequence Mapping Analysis

To understand the representation of our 18S rRNA gene-based OTUs in previous metabarcoding-based studies, we used a comprehensive amplicon sequence data set resulting from the merging of the EukBank V4 region amplicon dataset composed of 12,672 samples (Berney et al. 2023) and the V4 region amplicon subset of the MetaPR2 database v3.0.1 composed of 6,069 samples (Vaulot, et al. 2022). Representative sequences from 418,231 amplicon sequence variants (ASVs) were dereplicated using vsearch --derep_fulllength producing 386,482 unique sequences which were clustered at 97% identity using vsearch --cluster_fast --id 0.97 resulting in 166,372 short amplicon OTUs (short OTUs). We used the bbmap v39.33 (https://sourceforge.net/projects/bbmap/) software to map the short OTU representative sequences to the 13,238 metagenomic assembly OTU representative sequences using the following options: maxindel=2000, idfilter=0.80, ambiguous=best, and slow=t allowing sequence variation to account for the limitations of using short sequences of the V4 region and performing a sensitive search. The relationship between reference sequences and aligned amplicon sequences was obtained using the bamtobed option in BEDTools v2.31 (Quinlan & Hall, 2010). To get a measure of the diversity of sequence variants captured by each metagenomic OTU in the sequence mapping we used the weighted environmental abundance calculated using the following expression:

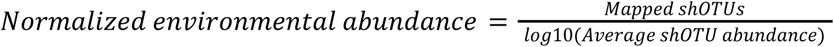

Short OTU (shOTU) abundance was obtained from the information provided in the EukBank and MetaPR2 datasets and used to normalize the number of aligned sequences to each OTU by the abundance of each short OTU as more abundant short OTUs are expected to be more likely to be found and aligned to a metagenomic OTU. To inspect the relationship between normalized environmental abundance and predicted primer binding bias, we calculated the Spearman correlation coefficient between the primer binding bias score and normalized environmental abundance using the spearmanr function from scipy v 1.16 (Virtanen et al. 2020).

### Correlation Between SSU rRNA Sequence Read Depth

For each of the most frequently represented ecosystem types in our dataset (Soil, Freshwater, and Marine), we evaluated the sequences within each 18S rRNA gene cluster, retaining only those with a minimum of three sequences across different samples from specific environments, resulting in the acquisition of ecosystem-filtered 18S rRNA gene clusters. An analogous procedure was conducted to retrieve ecosystem-filtered 16S clusters, from which we calculated Spearman’s rank correlation coefficients for each pair of 18S and 16S rRNA gene filtered clusters whenever a minimum of three samples contained sequences from both clusters. Adjustments for p-values were made using the Benjamini-Hochberg method for each GOLD ecosystem type independently, with a replication of this process for correlations among 18S rRNA gene sequence clusters, incorporating the additional step of excluding autocorrelations. Correlation coefficients and p-values are available in the supplementary data repository.

## Acknowledgments

The work conducted by the U.S. Department of Energy Joint Genome Institute (https://ror.org/04xm1d337), a DOE Office of Science User Facility, is supported by the Office of Science of the U.S. Department of Energy operated under Contract No. DE-AC02-05CH11231.

## Data availability

Source Data are provided with this paper. Sequence data including SSU rRNA sequences and SAG assemblies, together with tables containing the full metadata from sequencing datasets from IMGM, rarefied counts, correlations, tree files and sequence cluster information are available as a supplementary dataset via Zenodo with the following DOI: 10.5281/zenodo.20653200.

## Code availability

The pipeline for extracting long SSU rRNA sequences from metagenomic contigs developed for this study is available at https://github.com/NeLLi-team/ssuextract.

## Supplementary Figures

**Supplementary figure 1.**
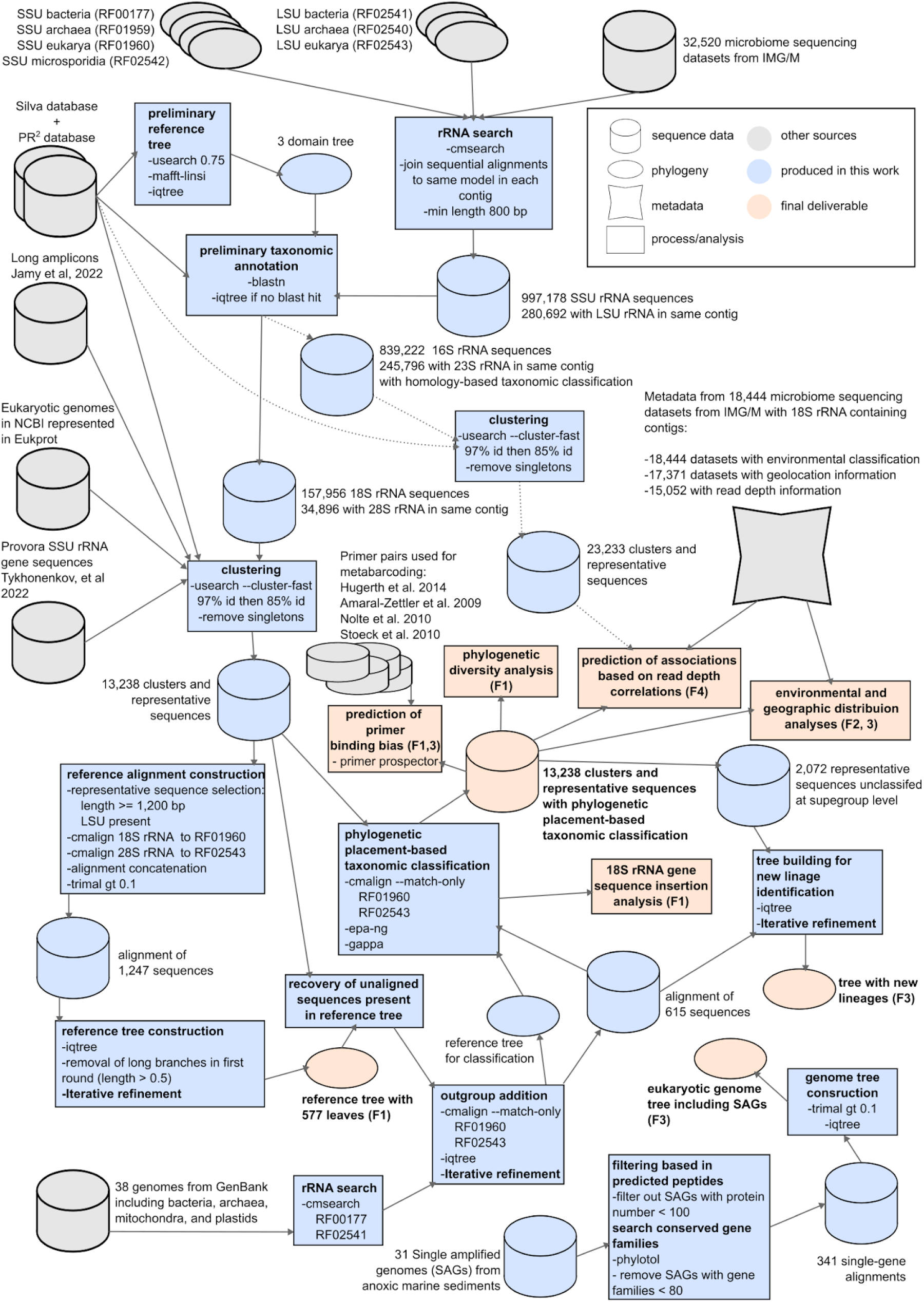
Summary of the methods used in this work. Cylinders correspond to sequence data, while the ovals indicate phylogenies, stars indicate metadata, and squares represent analyses. Some final deliverables include an F and one or two numbers in parenthesis, which indicate the figure of the main text in which they are included. “Iterative refinement” refers to the process of manually inspecting the tree and removing sequences representing rogue taxa and long branches that could produce artifacts in the topology from the seed alignment and rebuilding the tree.

**Supplementary figure 2.**
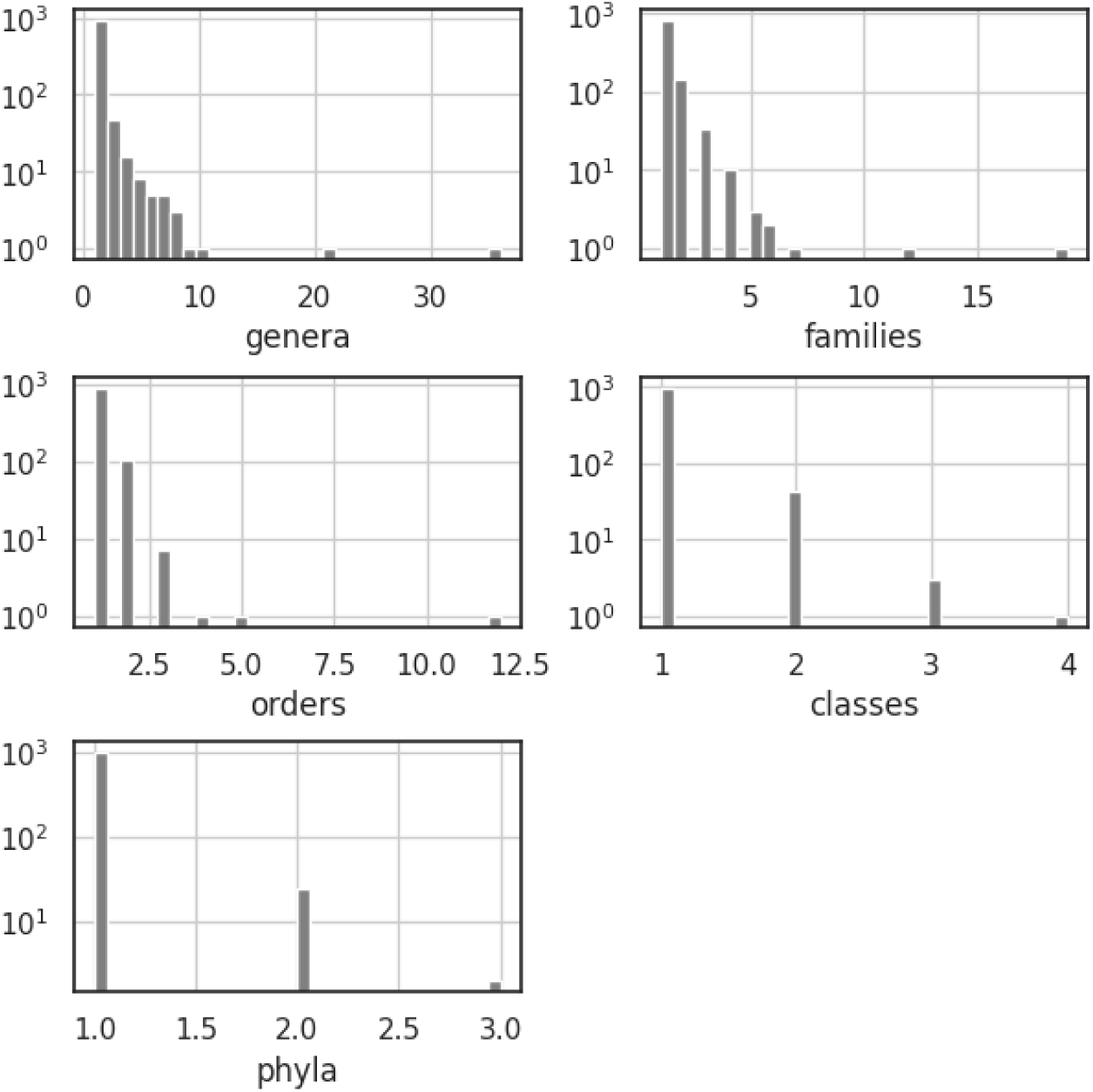
Distribution of clusters comprising any given number of different taxa within a single reference sequence-only cluster.

**Supplementary figure 3.**
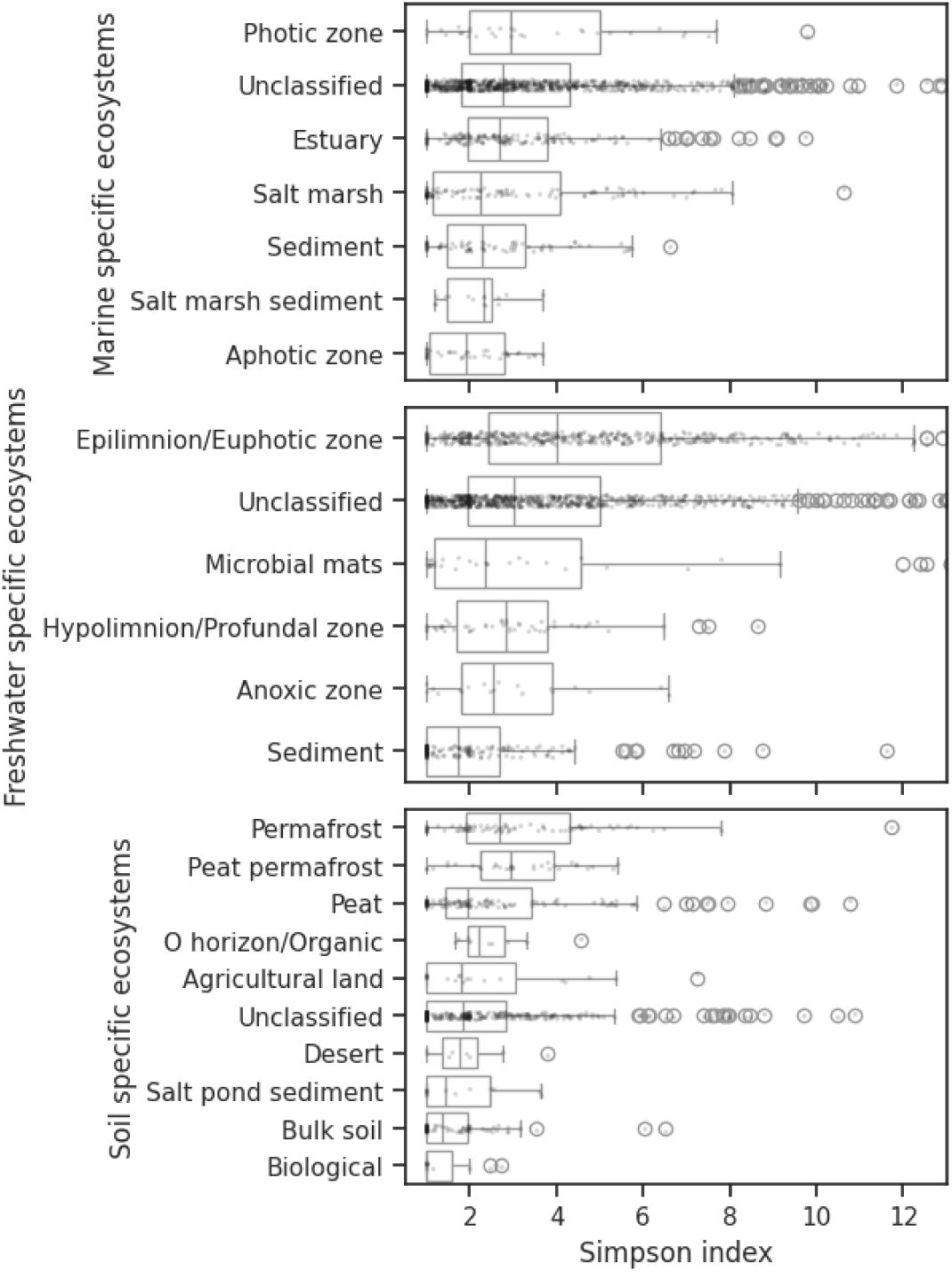
Per-sample diversity distribution for each specific ecosystem with at least 1% share of the datasets separated by ecosystem types. Box plots display medians (vertical lines), interquartile ranges (boxes), and data distribution (whiskers), with outliers shown as circles.

**Supplementary figure 4.**
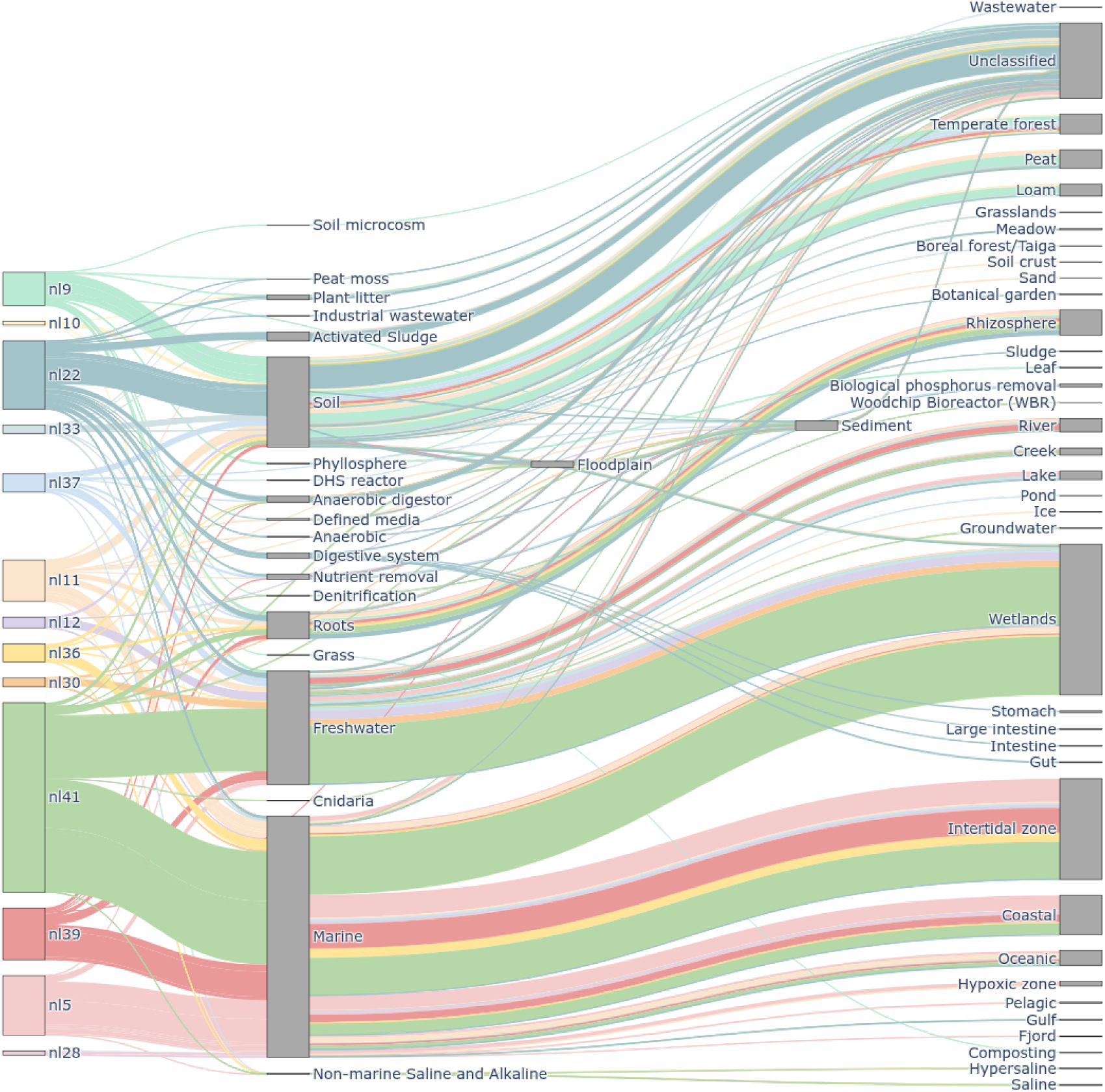
Contribution of each ecosystem subtype and ecosystem type to the observed pool of 18S rRNA gene sequences attributable to the 13 new lineages.\

**Supplementary figure 5.**
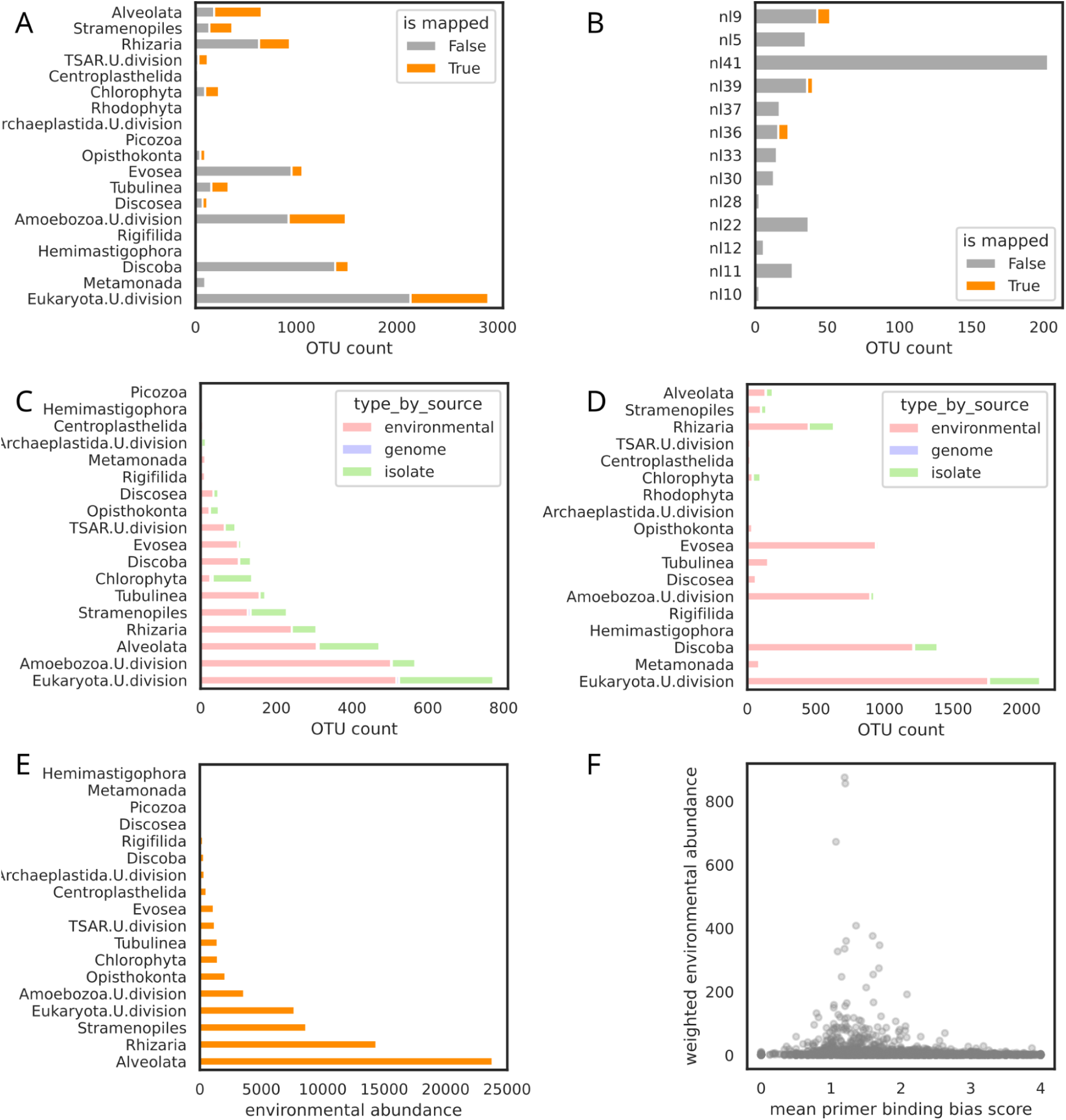
18S rRNA gene V4 region short amplicon sequence mapping to long metagenomic assembly OTU representative sequences. A. Number of metagenomic OTU representatives for each protist division colored by their alignment status to short amplicon OTU (shOTU) representatives. B. Number of metagenomic OTU representatives for each of the new lineages presented in this work colored by their alignment status to shOTU representative sequences. C. Number of metagenomic OTU representatives that were aligned at least once with shOTU representative sequences colored by the cluster type regarding the source of the sequences belonging to each OTU. D. Number of metagenomic OTU representatives that were not aligned at least once with shOTU representative sequences colored by the cluster type. E. Environmental abundance for each protist division expressed as the total number of alignments of metagenomic OTU representative sequences and shOTU representatives. F. Relationship between mean primer binding bias score and the weighted environmental abundance for each metagenomic OTU with alignments to short read sequences.

**Supplementary figure 6.**
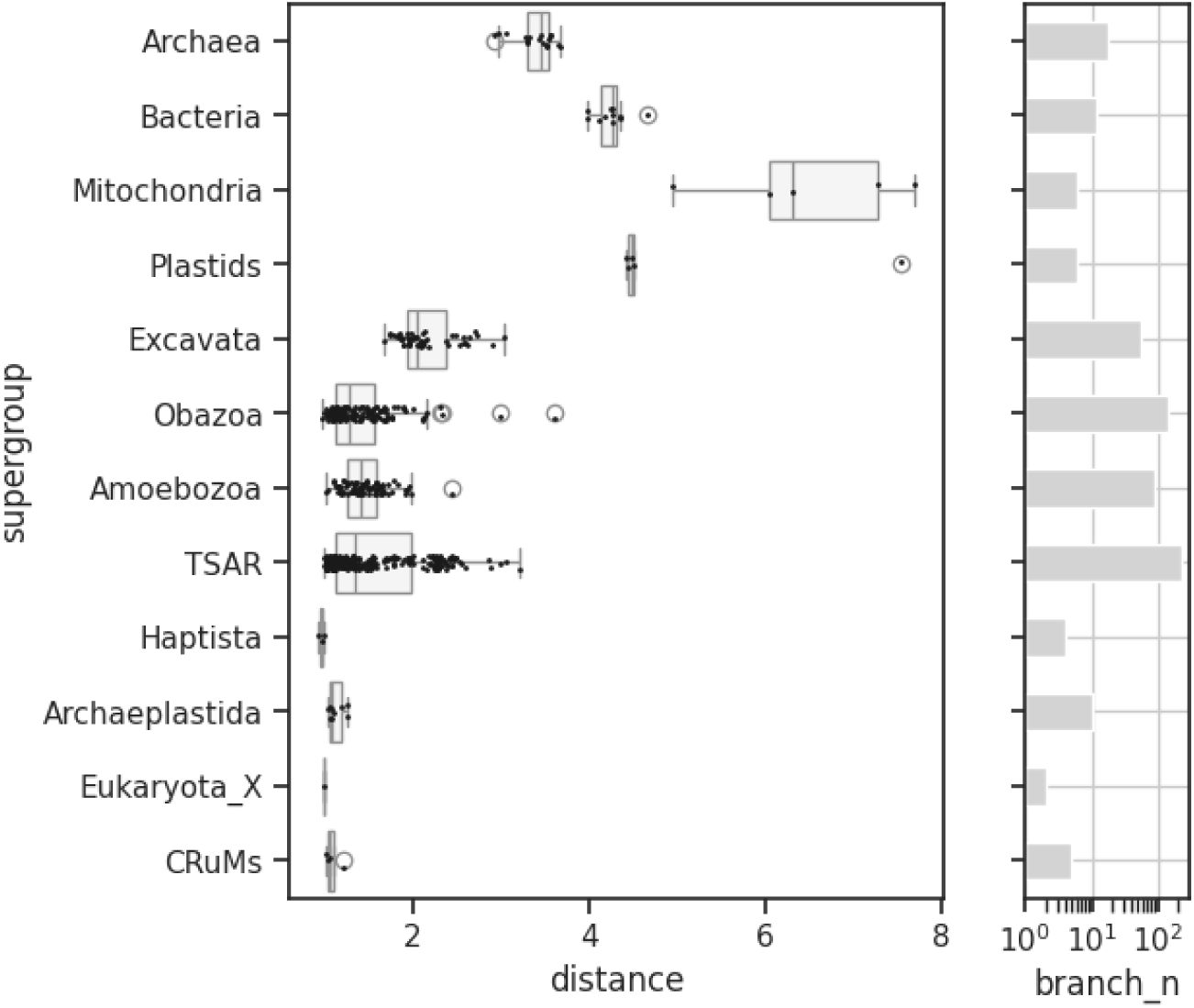
Tree distance between branches belonging to NL11 and other taxonomic major groups. Left: Distribution of distances between the lowest common ancestor of NL11 and branches belonging to each eukaryotic supergroup and prokaryotic groups. Box plots display medians (vertical lines), interquartile ranges (boxes), and data distribution (whiskers), with outliers shown as circles. Right: number of branches belonging to each taxonomic category in log10 scale.

**Supplementary figure 7.**
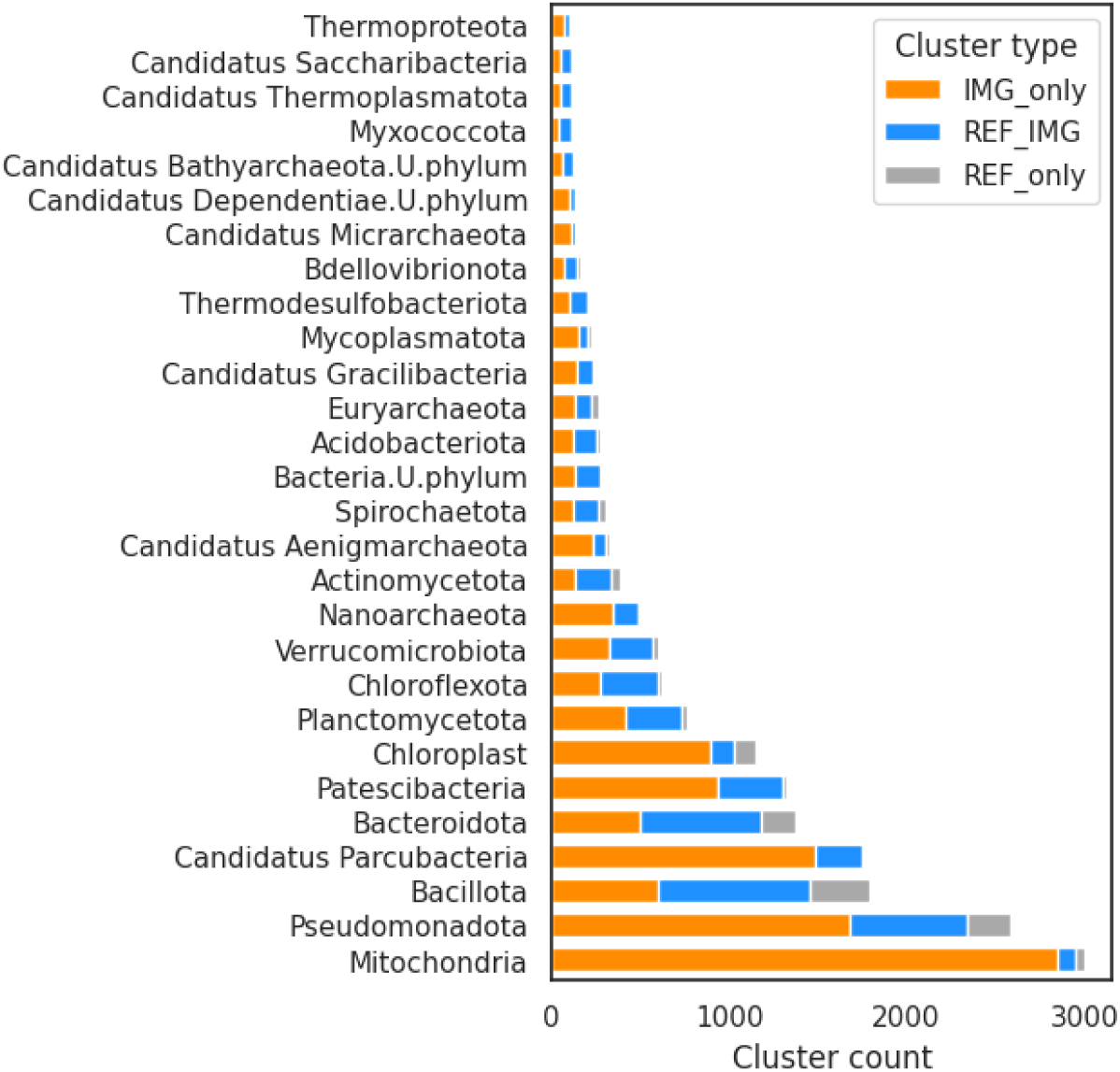
Abundance of clusters for each prokaryotic taxonomic group representing at least 0.5% of the total number of clusters within the analyzed environmental samples. Bars are partitioned according to their share of cluster types as defined by the origin of the member sequences. REF_IMG: includes both sequences from IMG and from reference databases; REF_only: includes sequences from reference databases exclusively.

## Supplementary Tables

Supplementary Table 1. Number of sequencing datasets with observed protist 18S rRNA sequences by Ecosystem Subtype.

Supplementary Table 2. Counts of samples with no observed protist 18S rRNA sequences by Ecosystem Subtype filtered by Ecosystem Subtype and Specific Ecosystem.

Supplementary Table 3. Mean prevalence, abundance and weighted abundance data of protist 18S rRNA sequences among ecosystem types at the family level.

Supplementary Table 4. Metadata and frequency of 18S rRNA genes for samples in which new lineages were found.

Supplementary Table 5. Functional and taxonomic annotations for predicted proteins belonging to contigs extracted using tax-aliquots on sequences belonging to NL5.

Supplementary Table 6. Functional and taxonomic annotations for predicted proteins belonging to contigs extracted using tax-aliquots on sequences belonging to NL11.

Supplementary Table 7. Kruskal-Wallis test results and paired two-tailed Mann-Whitney results for primer binding bias scores between protist divisions.

Supplementary Table 8. Information on selected genomes of archaea, bacteria, mitochondria and plastids used as outgroup for the tree built for phylogenetic placement-based taxonomic classification.

Supplementary Table 9. Kruskal-Wallis test results and paired two-tailed Mann-Whitney results for insertion counts in the 18S rRNA gene between protist divisions.

Supplementary Table 10. Kruskal-Wallis test results and paired two-tailed Mann-Whitney results for insertion lengths in the 18S rRNA gene between protist divisions.

Supplementary Table 11. Estimated model parameters and statistics for OTU accumulation curves by Ecosystem Type.

Supplementary Table 12. Kruskal-Wallis test results and paired two-tailed Mann-Whitney results for primer binding bias scores between 18S OTU categories defined by their affiliation to new lineages, source type and presence in the 18S-28S reference tree.

Supplementary Table 13. Kruskal-Wallis test results and paired two-tailed Mann-Whitney results for tree distances between NL11 and taxonomic groups.

## Supplementary Methods

To search for rRNA sequences taking into account the extreme variations in length due to introns and other additional sequences in eukaryotic genomes, we developed a workflow named “ssuextract”. This workflow uses the Infernal software (Nawrocki and Eddy 2013) to search a covariance model (CM) to nucleic acid sequencing contigs and search homologs using blastn (Camacho et al. 2009) against the Silva (Quast et al. 2013) and PR2 (Guillou et al. 2013) databases as a first approach to get a taxonomic classification for each sequence. It was designed with the 18S rRNA gene in mind, but it can also be used to extract 28S, 16S, 23S rRNA gene sequences, and practically any nucleic acid sequence for which a covariance model is available.

The key component of this workflow is the detection of long sequences represented by the covariance model through multiple sequential hits along a single contig reported in the cmsearch output table produced using the –anytrunc option (i.e. a fragmented alignment). Full alignments to the model in which the start and end position of the covariance model are found within the subject contig are returned instantly, while repetitions of the covariance model in the contig (i.e. paralog sequences) are also considered and extracted individually. This process is implemented in Python3 as illustrated with the following block of pseudocode:

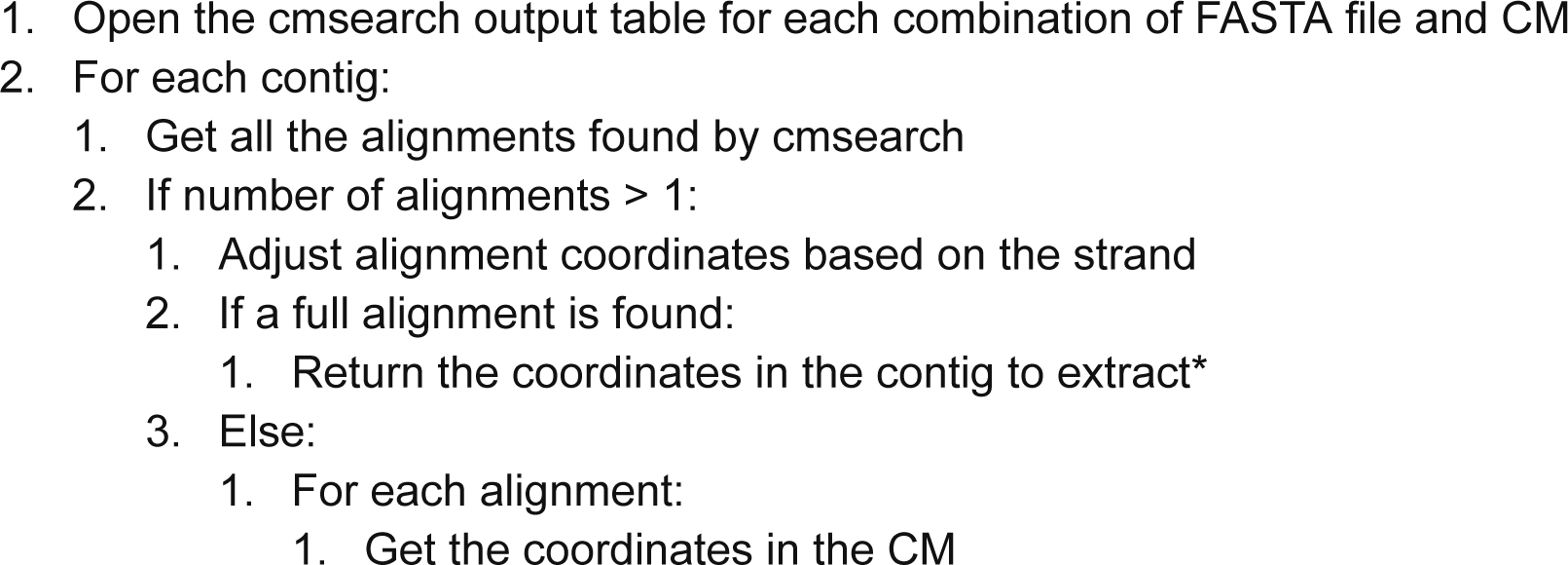
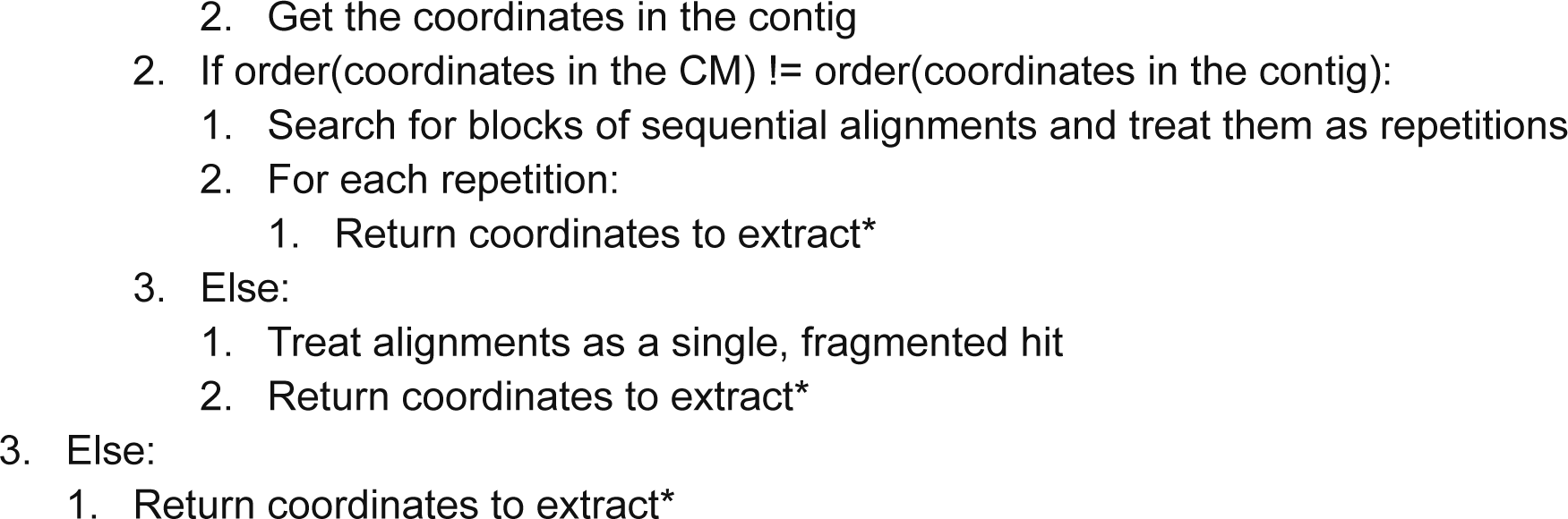

Asterisks denote the different final outcomes for the alignment information on each contig. Only sequences representing an alignment to the CM with length >= 800 bp are extracted for further analyses. The workflow uses snakemake (Mölder et al. 2021) to manage sub processes in an ordered and scalable manner, and anaconda (https://anaconda.com/) to install and load dependencies such as Infernal and Blast+. The code is publicly available at GitHub (https://github.com/NeLLi-team/ssuextract) and has been successfully tested in SUSE Linux Enterprise Server 15 SP5 and Ubuntu 24.04 LTS in an WSL instance for Windows 11.

